# Grubraw, a chemogenetic mitochondrial activator, reveals new mechanisms underlying the Warburg effect

**DOI:** 10.1101/2023.04.18.537329

**Authors:** Ekaterina S. Potekhina, Dina Y. Bass, Alexander V. Ivanenko, Alexander A. Moshchenko, Dmitri A. Korzhenevskiy, Evgeniy Shevchenko, Anastasia E. Karnaeva, Natalia F. Zakirova, Alexander V. Ivanov, Liubov E. Shimolina, Marina V. Shirmanova, Olga V. Lyang, Olga I. Patsap, Olga M. Kudryashova, Guzel R. Gazizova, Elena I. Shagimardanova, Oleg A. Gusev, Ivan Bogeski, Alexey M. Nesterenko, Vsevolod V. Belousov

**Author notes:** **Author Information** Correspondence and requests for materials should be addressed to E.S.P., or V.V.B. These authors contributed equally to this work (Ekaterina S. Potekhina, Dina Y. Bass).

## Abstract

Upregulation of glycolysis and downregulation of mitochondrial oxidative phosphorylation, termed as the Warburg effect, are characteristic of tumor cells^1,2^. Restriction of pyruvate flux into the mitochondrial matrix is one of the major mechanisms underlying this phenomenon^3^. Warburg-type metabolism is beneficial for rapidly proliferating cells, however its function remains unclear. Moreover, it is unknown what the metabolic consequences of activation of mitochondrial respiration in Warburg-type cancer cells are. Here we created a chemogenetic instrument, Grubraw, that generates pyruvate directly in the mitochondrial matrix bypassing restricted pyruvate influx. In cancer cells, Grubraw-driven pyruvate synthesis in the matrix increased mitochondrial membrane potential, oxygen consumption rate, and the amounts of TCA cycle intermediates. In a mouse model of human melanoma xenografts, chemogenetic activation of mitochondria caused a decrease in tumor growth rate. Surprisingly, cancer cells actively exported pyruvate generated by Grubraw in the mitochondria into the extracellular medium. In addition, activation of mitochondria induced downregulation of transcription of the genes that drive cell cycle progression, cell proliferation and DNA replication. Our results demonstrate that cells with Warburg-type metabolism use previously unknown mechanisms of carbon flux control to dispose of excessive mitochondrial pyruvate, and activation of mitochondria in these cells downregulates cellular proliferation.

## Main

Pyruvate is one of the key molecules critical to eukaryotic metabolism as the endpoint of glycolysis and an important branching point between lactic fermentation and oxidative phosphorylation (OXPHOS). In cancer cells, pyruvate is one of the keystone molecules for metabolic reprogramming, termed the Warburg effect. Current understanding describes the Warburg phenotype as metabolic reprogramming associated with upregulation of glycolysis and downregulation of OXPHOS^1,2^. Carcinogenesis and the Warburg effect are frequently accompanied by decreased pyruvate entry into mitochondria via mitochondrial pyruvate carrier (MPC)^3,4^. Moreover, pyruvate metabolism plays a significant role in tumor angiogenesis^5^, epithelial-mesenchymal transition (EMT), the development of chemoresistance in cancer cells^6^, and HIF-1α degradation inhibition under aerobic conditions^7^. Why cancer cells restrict mitochondrial function, and what the metabolic and phenotypic consequences are if mitochondria in cancer cells are forced to metabolize pyruvate and respire intensively remains unknown.

In order to force mitochondria in cancer cells to metabolize pyruvate and respire more intensively we have developed a chemogenetic molecular tool based on *Pseudomonas aeruginosa* FAD-dependent D-amino acid dehydrogenase DadA that allows bypass of the pyruvate transport block and activation of mitochondrial respiration by generating pyruvate from D-alanine directly in the mitochondria of eukaryotic cells and donating electrons downstream of Complex I. DadA-dependent oxidative deamination of D-alanine leads to production of pyruvate and ammonia, involving an unknown electron acceptor for FADH_2_ (Fig. 1a)^8^. Since D-alanine is absent in most mammalian tissues, except the brain and pancreas^9,10^, DadA activity in the cells can be regulated by external administration of D-alanine.

**Figure 1.**
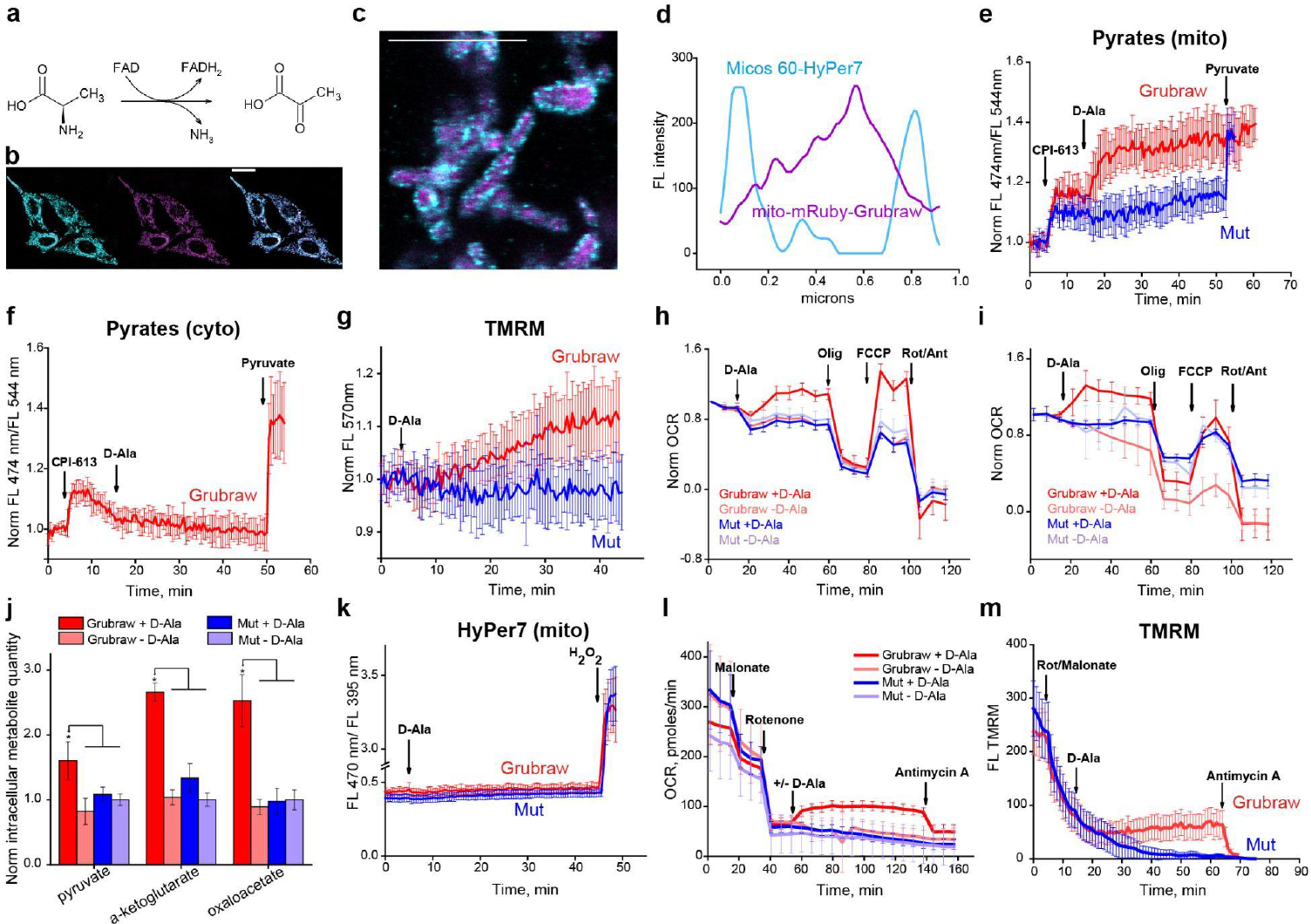
Grubraw activity in the mitochondria of the Hela Kyoto cells activates OXPHOS. **a**, The conversion of D-alanine to pyruvate catalyzed by DadA. **b**, Fluorescence image of HeLa Kyoto cells expressing mito-mRuby-DadA stained with mitoTracker Green. **c**, Confocal fluorescence image of HeLa Kyoto cells expressing mito-mRuby-DadA and Micos60-HyPer7. **d**, A representative fluorescence intensity profile along a cross-section of a mitochondrion from **c. e**, Mitochondrial pyruvate concentration in the presence of 120 μM CPI-613 by Pyrates sensor localized in the mitochondria. n = 14 – 15. **f**, Cytosolic pyruvate concentration in the presence of 120 μM CPI-613 by Pyrates sensor localized in the cytosol. n = 6. **g**, Mitochondrial membrane potential by TMRM. n = 12 – 23. **h**, The effect of Grubraw activity on OCR in HBSS medium without glucose. n=5. **i**, The effect of Grubraw activity on OCR in DMEM medium with 5 mM glucose. n=5. **j**, The effect of Grubraw activity on intracellular TCA cycle metabolites. Mean normalized values ±SD are shown. **k**, ROS production by HyPer 7 sensor localized in the mitochondria. n = 8. **l**, The effect of Grubraw activity on OCR in HBSS medium without glucose, in presence of 5 mM malonate, 30 μM rotenone, and 10 μM antimycin A. n=5. **m**, Mitochondrial membrane potential by TMRM in presence of 30 μM rotenone, 5 mM malonate, and 10 μM antimycin A. n = 15. In all experiments, where indicated, 10 mM D-Ala was used. Scale bar represents 25 μm in **b** and 5 μm in **c**. In **e, f, k, l**, red lines are for HeLa Kyoto cells expressing mRuby-Grubraw and blue lines for the cells expressing mRuby-Grubraw-mut. In **g, m**, red lines are for HeLa Kyoto cells expressing EGFP-Grubraw and blue lines for the cells expressing EGFP-Grubraw-mut. In **e, f, g, h, i, k, l, m**, mean or mean normalized values ±SD for n cells or wells are shown for every time point.

The fusion proteins mRuby-DadA and EGFP-DadA, retaining near-native enzymatic activity, were created to enable DadA tracking in live cells (Extended Data Fig. 1a). To control the unspecific effects of heterologous protein expression, an inactive mutant was created by changing the conservative Tyr17 and Tyr18 residues in the predicted FAD-binding domain to Ala residues (Extended Data Fig. 1b, 1a).

The fusion proteins were targeted to the mitochondrial matrix, and mitochondrial localization was confirmed by imaging with the mitochondria-specific dyes TMRM or MitoTracker Green (Fig. 1b, Extended Data Fig. 1c, 1d). Mito-mRuby-DadA matrix localization in HeLa Kyoto was confirmed by coexpressing with Micos60-HyPer7 protein localized in the mitochondrial intermembrane space (Fig. 1c). The fluorescence intensity profile along the mitochondrion cross-section showed that the mRuby fluorescence peak was located between two HyPer7 fluorescence peaks, confirming that all observed DadA was localized in the mitochondrial matrix (Fig. 1d). Since the purpose of our study was to reverse the Warburg effect, we named the mitochondrial DadA version fused to the fluorescent protein **Grubraw** (reversed Warburg). To directly monitor pyruvate dynamics in the cytosol and mitochondrial matrix of HeLa cells expressing Grubraw, we used Pyrates^11^, a genetically encoded fluorescent sensor for pyruvate. To prove that Grubraw generates pyruvate in the matrix when supplied with the substrate, we removed glucose from the cell culture medium, and inhibited pyruvate entrance to the TCA cycle using the inhibitor of pyruvate dehydrogenase CPI-613. Upon D-Ala addition, we observed a significant increase in matrix pyruvate concentration (Fig. 1e). Grubraw activity did not affect the concentration of pyruvate in the cytosol (Fig. 1f) suggesting that HeLa cells maintain a steady pyruvate concentration in this compartment.

We were also interested in whether loading the mitochondrial matrix with pyruvate could enhance electron transport chain (ETC) function. Indeed, Grubraw activity increased mitochondrial transmembrane potential in HeLa cells (Fig. 1g) indicating that the products of Grubraw activity (pyruvate and FADH_2_) enter mitochondrial oxidative metabolism. D-Ala itself did not affect mitochondrial transmembrane potential.

Moreover, Grubraw activity increased oxygen consumption rate in HeLa cells both under fasting conditions (Fig. 1h) and in rich culture medium (Fig. 1i). We concluded that in HeLa cells, even when substrates in the medium are sufficient, the ETC does not function at the maximal physiological rate, likely due to the regulation of mitochondrial transporters and other metabolic “switching nodes” biased toward aerobic glycolysis. Pyruvate and/or FADH_2_ produced by Grubraw can contribute to ETC function independently of these mechanisms and increase the proportion of OXPHOS in cellular metabolism.

What is more, Grubraw activity increased the intracellular concentrations of some TCA cycle metabolites, namely α-ketoglutarate and oxaloacetate, as shown by mass-spectrometry (Fig. 1j). Moreover, intracellular concentration of pyruvate also increased. These results suggest that not only was ETC function affected by Grubraw activity, but general matrix metabolism was also enhanced, likely due to the entrance of Grubraw-derived pyruvate into the TCA cycle. Thus, the increase in the matrix pyruvate concentration *per se* is able to enhance the TCA cycle flux.

Since we had no direct data about the acceptor of electrons from FADH_2_ of Grubraw, it was essential to confirm that they are not transferred to O_2_ thus generating ROS, and that Grubraw activity does not induce oxidative stress in mitochondria. Moreover, the Warburg effect is sometimes thought to be a part of the antioxidant defense system in cancer cells^2^. Given these considerations, we used the ultrasensitive genetically encoded H_2_O_2_ probe HyPer7 targeted to the mitochondrial matrix^12^. No changes in HyPer7-mito signal were detected after D-Ala addition in HeLa cells expressing active and inactive Grubraw (Fig. 1k). Thus, Grubraw does not generate physiologically significant H_2_O_2_ concentrations^13^, nor does acceleration of OXPHOS elevate endogenous mitochondrial H_2_O_2_ production.

For the DadA homologue from *Esherichia coli*, electron transfer from FAD to Q_10_ analogues with various isoprenoid side chain lengths was demonstrated ^14^. Since Grubraw was active in mitochondria and did not generate ROS, and since the purified enzyme was not able to reduce NAD^+^ or NADP^+^ *in vitro* (data not shown), we speculated that it could transfer its electrons to some specific acceptor in the ETC, most probably to Q_10_. In presence of the complex I inhibitor rotenone and complex II inhibitor malonate, Grubraw activity prevented complete loss of the mitochondrial transmembrane potential and oxygen consumption, in contrast to the inactive control enzyme version (Fig. 1l, 1m). After the addition of complex III inhibitor antimycin A, mitochondrial transmembrane potential and oxygen consumption decreased even in the cells with active Grubraw. This suggested that the electrons from Grubraw entered the ETC downstream of complexes I and II and upstream of complex III, likely via the Q-pool, and thus partially maintained mitochondrial function. Additionally, rotenone and all other small-molecule inhibitors used in this work were proven *in vitro* to have no effect on Grubraw enzymatic activity (Extended Data Fig. 1e).

Moreover, the genetically encoded pH sensor SypHer3s^15^ with a matrix localization sequence did not report any significant pH changes when Grubraw was active (Extended Data Fig. 1f). Thus, ammonia released during Grubraw functioning did not affect mitochondrial pH.

Grubraw activity significantly reduced the growth rate of HeLa cells, but D-Ala itself did not affect proliferation. (Extended Data Fig. 1g). These data suggest that oxidative metabolism intensification, at least at the level induced by Grubraw, significantly alters cellular proliferation. The increase in TCA cycle metabolite pools generated by Grubraw could affect not only the mitochondrial oxidative reactions, but also processes in the cytosol. For example, the increase in oxaloacetate and α-ketoglutarate pools might enhance the function of the malate-aspartate shuttle, which in turn might change the cytosolic NAD^+^/NADH ratio. Indeed, the genetically encoded fluorescent sensor SoNar^16^ demonstrated oxidation of the cytosolic NAD pool during Grubraw activity in HeLa cells (Fig. 2a).

**Figure 2.**
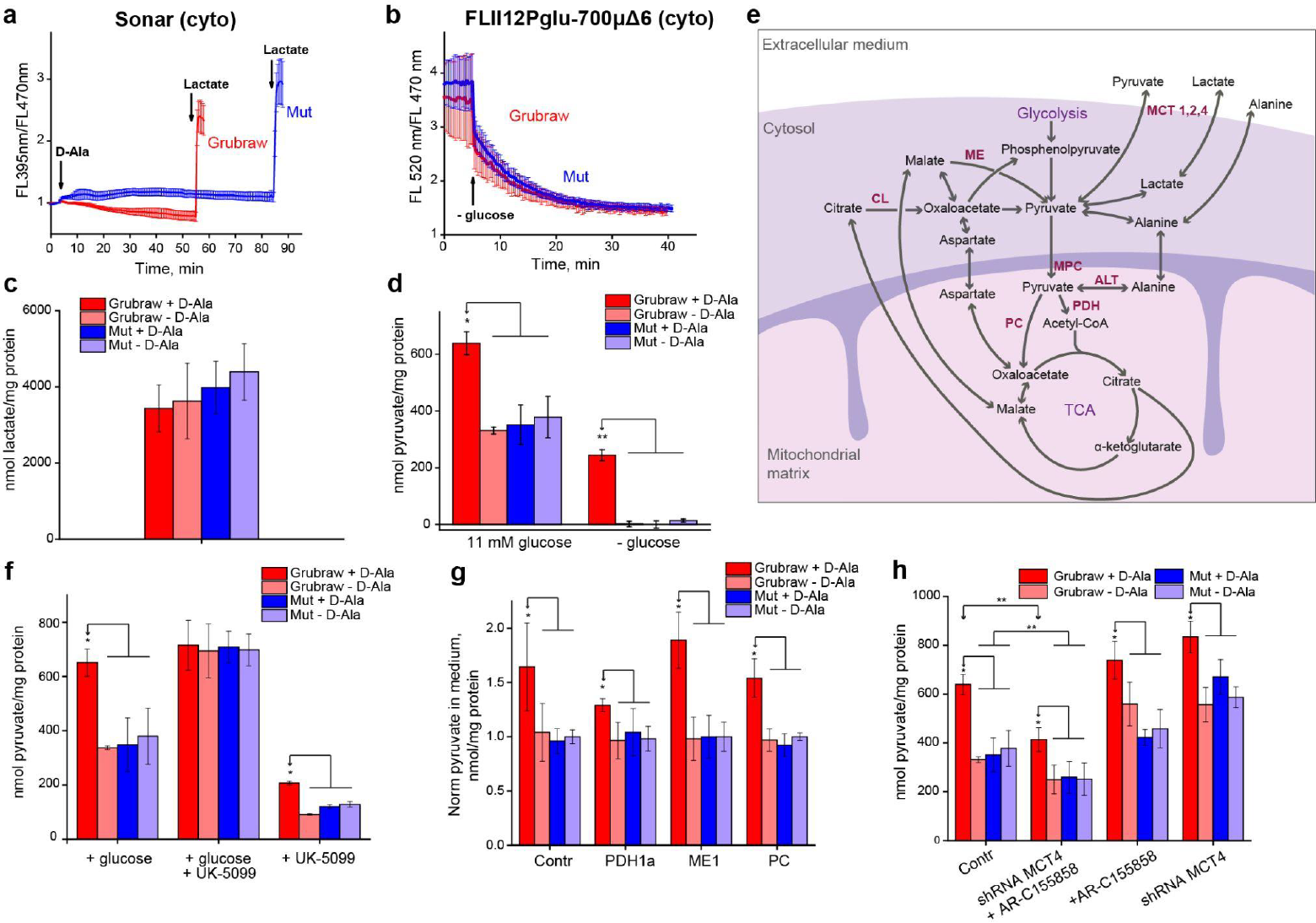
Grubraw activity increases extracellular pyruvate export in HeLa cells|. **a**, NAD+/NADH ratio by SoNar sensor. n = 14 – 18 cells. **b**, Glycolysis rate by FLII12Pglu-700μΔ6 glucose sensor. n = 14 cells. **c**, The effect of Grubraw activity on extracellular lactate. **d**, The effect of Grubraw activity on extracellular pyruvate in presence and absence of 11 mM glucose. **e**, Possible biochemical pathways for export of pyruvate from the mitochondrial matrix into the extracellular medium. The key enzymes and transporters are labeled in purple letters, and the metabolites are labeled in black letters. **f**, The effect of Grubraw activity on extracellular pyruvate. MPC activity was inhibited by 10 μM UK-5099 in presence and absence of 11 mM glucose. **g**,.The effect of Grubraw activity on extracellular pyruvate. PDH1a, ME1, and PC were downregulated by RNA interference. The control cells expressed a scrambled shRNA. In all samples, 11 mM glucose was present in the medium. **h**, The effect of Grubraw activity on extracellular pyruvate. MCT 1,2 and/or MCT4 activity was inhibited using 0.5 μM AR-C155858 and RNA interference, respectively. In all samples, 11 mM glucose was present in the medium. In all experiments, where indicated, 10 mM D-Ala was used. In **a** and **b**, red lines are for HeLa Kyoto cells expressing mRuby-Grubraw and blue lines are for cells expressing mRuby-Grubraw-mut. Mean or mean normalized values ±SD for n cells are shown at every time point. The data in **c**, **d**, **f**, **h** represent mean pyruvate or lactate quantity in the medium normalized to protein quantity in the sample ±SD. In plot **g**, these values were normalized to the corresponding value in the ″Grubraw-mut - D-alanine″ sample. Mean values ±SD are shown. *p<0.05

The increase in cytosolic NAD^+^ concentration might increase the rate of glycolysis, since the NAD^+^ supply is a limiting factor in this pathway, especially in cells of the Warburg metabolic type. However, the Grubraw activity did not change the rate of intracellular glucose metabolization, as shown by the genetically encoded fluorescent sensor for glucose FLII12Pglu-700μΔ6^17^ (Fig. 2b).

The cytosolic NAD^+^/NADH ratio is coupled to the balance of lactate and pyruvate production by LDH. Thus, we hypothesized that Grubraw-induced cytosolic NADH depletion results in decreased lactate export. However, no significant changes in extracellular lactate concentration were observed in HeLa cells expressing Grubraw after incubation with D-alanine compared to the control cells expressing Grubraw-mut and/or incubated without D-alanine (Fig. 2c).

Therefore, we speculated that the change in cytosolic NAD^+^/NADH ratio is compensated by enhanced pyruvate export, despite not being conventional for Warburg metabolic cells type. Indeed, Grubraw activity increased extracellular pyruvate concentration in HeLa cell culture (Fig. 2d) both in the presence and absence of glucose in the culture medium. It should be noted that, in the presence of glucose, some basal level of pyruvate export existed in all samples. Under fasting conditions, no extracellular pyruvate was detected in the samples before D-alanine addition. These data show that HeLa cells strictly regulate OXPHOS rate and matrix pyruvate concentration, exporting the excessive pyruvate into the extracellular medium.

We were next interested in the pathway by which pyruvate exits the mitochondrial matrix (Fig. 2e). MPC transports pyruvate molecules together with protons, so it is unlikely to perform reverse pyruvate transport against the proton concentration gradient. Indeed, incubation of the cells in glucose-free medium with the selective MPC inhibitor UK-5099 (10 μM) did not prevent enhanced Grubraw-dependent pyruvate export (Fig. 2f). In the control samples, however, pyruvate concentration also was not zero, probably because the residual cytosolic pyruvate could not be transported into mitochondria and was directed to the extracellular medium instead. It is of note that, UK-5099 did not influence the enzymatic activity of Grubraw *in vitro* (Extended Data Fig. 2b) nor D-alanine transport (Extended Data Fig. 1h). The latter result indicates that D-alanine transport into the mitochondrial matrix does not depend on MPC. We demonstrated that in presence of glucose, the extracellular pyruvate concentration increased in all control samples compared to the samples without UK-5099, probably because of the increased proportion of glycolysis-derived pyruvate in the export flux (Fig. 2f).

Therefore, we hypothesized that matrix pyruvate is converted into some intermediates which are transported to the cytosol and turned back into pyruvate there. These might be aspartate, malate, citrate, or alanine (Fig. 2e). To clarify which exact mechanism is used, we knocked down key enzymes involved in the possible export pathways, namely pyruvate dehydrogenase (PDH), pyruvate carboxylase (PC), and malic enzyme (ME1), in HeLa Kyoto cells expressing active and inactive versions of Grubraw. The mRNA quantities of PDH1a subunit, PC, and ME1 (cytosolic isoform) were decreased roughly 30-, 4-, and 5-fold, respectively, compared to the control cell line expressing scrambled shRNA (Extended Data Fig. 2a), which resulted in a significant protein quantity decrease, as shown by mass-spectrometry analysis (Extended Data Fig. 2b). Grubraw-dependent pyruvate export was retained in all knockdown cell lines. However, in the cells with PDH1a knockdown the difference was markedly decreased (Fig. 2g). This indicates that pyruvate is oxidized and enters the TCA cycle to be exported from the mitochondrial matrix.

Next, we studied the roles of monocarboxylate transporters (MCTs) in Grubraw-induced pyruvate export. MCTs 1, 2, 3, and 4 perform proton-coupled transport of pyruvate, lactate, and some other monocarboxylic acids through the plasma membrane. However, MCT3 is expressed only in some tissues of the eye^18^, so we focused our attention on MCTs 1, 2, and 4. We obtained HeLa Kyoto cell lines with stable active and inactive Grubraw expression and shRNA-mediated MCT4 knockdown (MCT4 mRNA quantity decreased roughly 9-fold, (Extended Data Fig. 2a and b). On the other hand, we inhibited MCT1 and MCT2 activity using the selective inhibitor AR-C155858 (0.5 μM). None of these conditions alone affected pyruvate export. However, when MCT4 knockdown and MCT1 and MCT2 inhibition were performed simultaneously, extracellular pyruvate concentrations proportionally decreased in all samples (Fig. 2h). These data suggest that MCT1, MCT2, and MCT4 are all involved in pyruvate export and are interchangeable.

To better understand how Grubraw-derived pyruvate enters cell metabolism and shifts metabolic fluxes, we performed ^13^C metabolic labeling experiments. HeLa cells expressing Grubraw were cultured for 24 hours in a medium with labeled carbon sources (glucose or glutamine). Focusing on acute effects of Grubraw activation, we changed the growth media to HBSS solution containing labeled carbon substrate (with only labeled glucose, or a mixture of glucose and glutamine with only one labeled component), added D-alanine and incubated cells under these conditions for 90 minutes before extraction (Extended Data Fig. 3d).

In labeled glucose only medium, we used 1,2-^13^C-glucose as it can provide information on glycolysis and the pentose-phosphate pathway^19–21^ (Fig. 3a, 3b). Intra- and extracellular pyruvate in this experiment had almost the same isotopologue distribution (Fig. 3c, Extended Data Fig. 3b) with about half of all pyruvate having two labeled carbons. Activation of Grubraw for 90 min led to a significant decrease in the M+2 isotopologue relative to M+0 in both intra- and extracellular pyruvate, intracellular α-KG and OA, and extracellular lactate (Fig. 3c). This “isotopic dilution” occurs because pyruvate from D-alanine evidently has a natural isotopic distribution, confirming the existence of a carbon flux from the Grubraw-derived mitochondrial pyruvate pool to the pyruvate pool in the extracellular medium.

**Figure 3.**
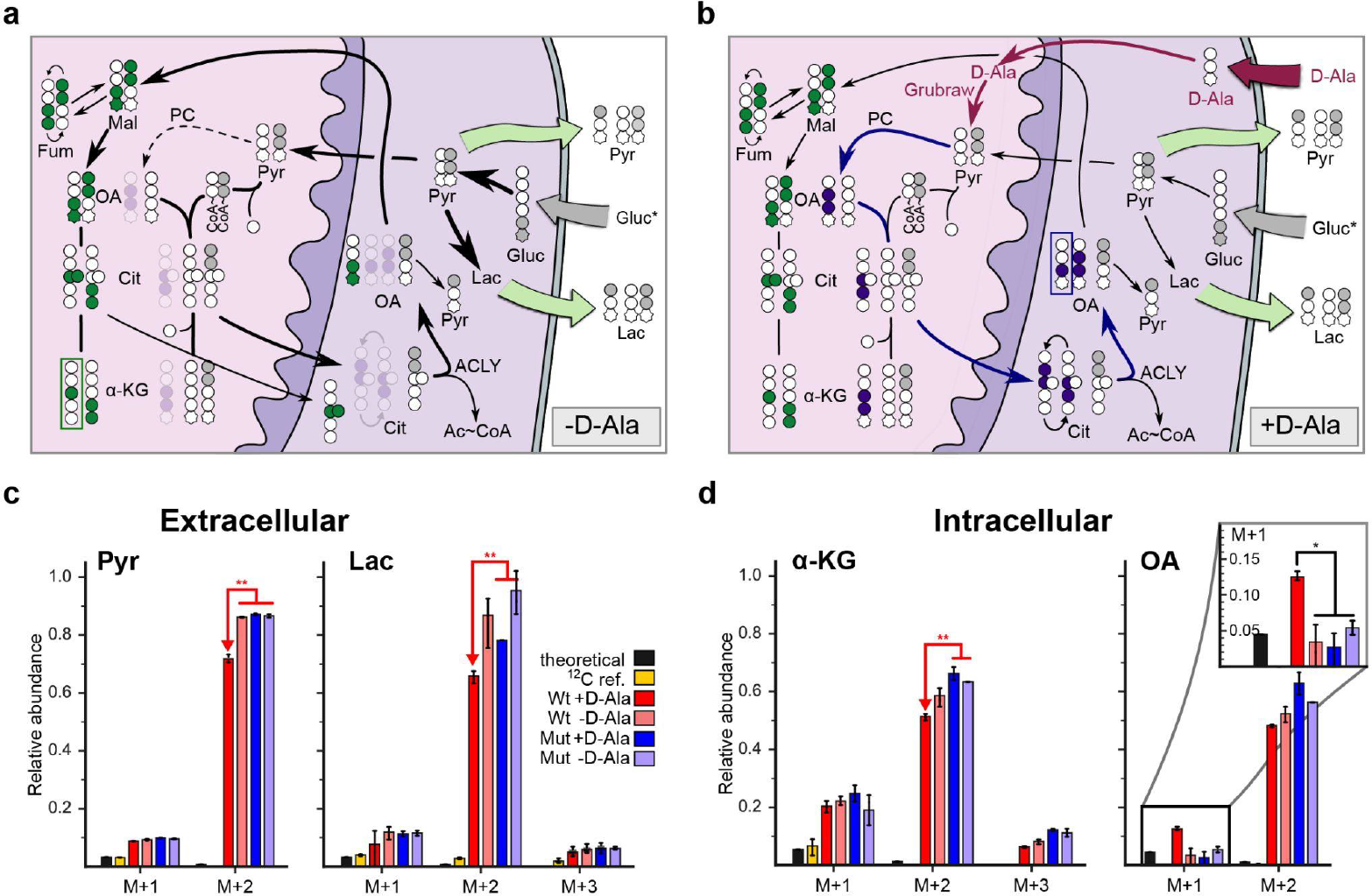
Major metabolic fluxes in HeLa cells growing on a medium with 1,2-^13^C_2_-glucose as the only carbon source. **a, b**, Carbon label migration from glucose over the central metabolism in the absence (**a**) and presence (**b**) of Grubraw activity. Labeled carbon atoms are colored gray 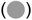 by default; green 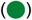 when they entered the matrix as the C4 metabolite; and blue 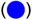 when they entered the TCA cycle via PC reaction. **c**, Isotopologue probability distribution for extracellular pyruvate and lactate molecules. **d**, Isotopologue probability distribution for intracellular OA and α-KG molecules. All probabilities are normalized to M+0 one. The derivatizer effect on the distributions was removed. Data on the labeling probability of C1 atoms for this experiment is provided in Extended Data Fig. 3a.

At the same time, the probability of OA M+1 behaves strictly differently (Fig. 3a, 3b). In all control cells, the probability of OA M+1 corresponds to the natural occurrence of the isotope, while the “Grubraw+D-Ala” sample contained an elevated OA M+1 isotopologue. In contrast, the probability of α-KG M+1 was increased in all labeled samples (Fig. 3d). Only the C1 α-KG carbon is removed during the [α-KG → OA] conversion in the TCA cycle. Thus, isotopologues with an increased content of α-KG M+1 must carry the tracer atom predominantly in the C1 position in order to turn into an OA pool with a negligibly small amount of tracer. However, fragmentation analysis (Extended Data Fig. 3a, 4) showed that only 10% of α-KG M+1 isotopologues were labeled at the C1 position. Therefore, we assume that OA is not synthesized from α-KG in TCA. This may be due to the incomplete TCA turning, which is characteristic of Warburg-type metabolism^21^. On the other hand, Grubraw-derived additional pyruvate influx can activate full turning of the TCA cycle and/or PC activity (Fig. 3b), resulting in an increase in M+1 OA isotopologue probability. Both activated metabolic pathways lead to OXPHOS intensification that correspond to our SeaHorse experiments (Fig. 1h, 1i).

We also conducted an experiment with two carbon sources, glucose and glutamine, considering these conditions to be closer to physiological conditions. Here we performed parallel labeling - the addition of two substrates, in which the first is labeled, and the second is unlabeled, and vice versa. We called these two states Q*G and G*Q for labeling with glutamine and glucose, respectively. Possible label migration through central metabolism from glutamine is shown on Fig. 4a and 4b, isotopomers distributions are shown on panels c – e.

**Figure 4.**
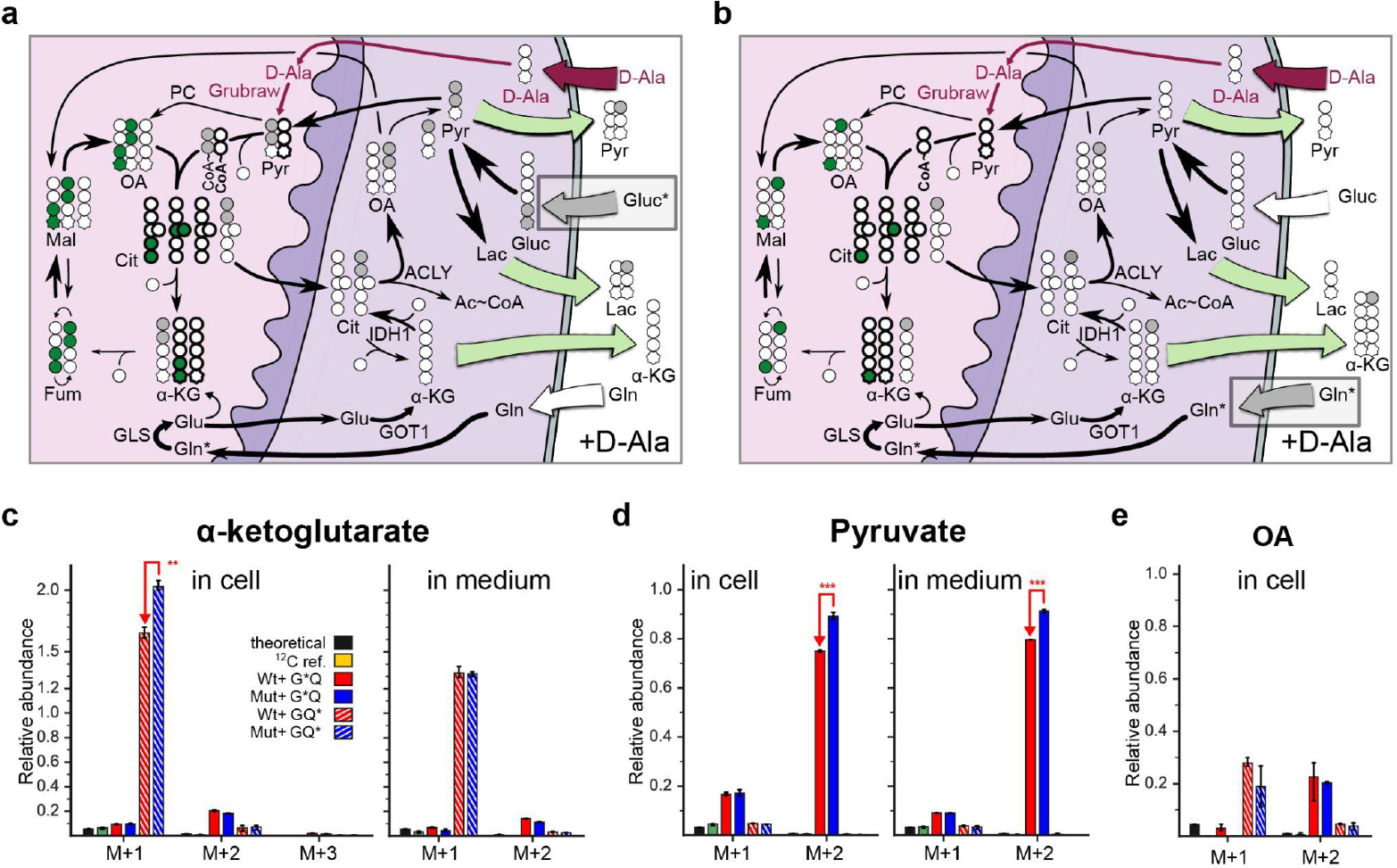
Major metabolic fluxes in HeLa cells growing on a poor medium with 1,2-^13^C-glucose/glutamine or glucose/5-^13^C-glutamine. **a, b**, Carbon label migration from glucose (**a**) and glutamine (**b**) in the presence of Grubraw activity. Labeled carbon atoms are colored gray 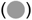 by default; green 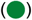 when they entered the matrix as the C4 metabolite. **c-e**, Isotopologue probability distribution of α-ketoglutarate (**c**), pyruvate (**d**) and oxaloacetate (**e**) Data for both intra- and extracellular metabolites are shown in (**c, d**). All probabilities are normalized to M+0. The derivatizer effect on the distributions was removed. Data on the labeling probability of the C_1_ atom for this experiment is provided in Extended Data Figure 3c.

Under these conditions, glucose supplied lower carbon input into the metabolism relative to glutamine: α-KG_M+2_ and OA_M+2_ relative abundance was about 20% in G*Q samples (Fig. 4c,e) versus roughly 60% in G* only samples (Fig. 3d). The influx of unlabeled carbons from D-Ala was detected by the significant dilution of Pyr_M+2_ in G*Q samples and intracellular α-KG_M+1_ in Q*G samples (Fig. 4d and 4c). Interestingly, dilution of α-KG_M+1_ by Grubraw activation in Q*G samples is observed in the intracellular pool, but not in the extracellular pool. One possible explanation is shown on Figure 4a and 4b. Glutamine-derived α-KG may enter the metabolism directly via the TCA cycle and indirectly via reductive carboxylation by IDH1 or IDH2. Glutamine catabolism begins with hydrolysis to glutamate catalyzed by matrix glutaminase. Subsequently, glutamate is deaminated to α-KG by aminotransferases in the cytosol or mitochondrial matrix (GOTx, GDH, AST). We propose that 5-carbon molecules can be transported from the mitochondrial matrix to the cytosol due to the concentration gradient, after which the glutamine tracer enters the metabolism predominantly via cytosolic IDH (IDH1):

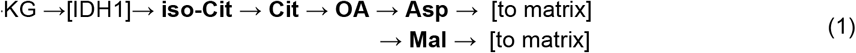

In this case, the α-KG carbon enters the matrix as the C4 metabolite after mixing with glucose-derived C4 metabolite. The cytosolic pool of α-KG does not mix directly with the mitochondrial pool and appears to be less sensitive to Grubraw-derived carbon influx. This hypothesis is also supported by the strong difference between the occurrence of α-KG_M+1_ and OA_M+1_ in Q*G samples: the abundance of α-KG_M+1_ is 200% relative to the M+0 isotopologue whereas for OA it is just 20%. If α-KG_M+1_ enters the matrix according to (1) under this condition, then 50% of OA becomes unlabelled. Further, OA is mixed in the matrix with OA obtained from glucose, and the probability of it being labeled drops to 16% (Fig. 4e).

In the experiments with the glutamine label, we did not see any tracer atoms in pyruvate. This is not surprising since pyruvate is formed primarily from glucose. The small part that could be derived from α-KG via pathway (2) is also mostly unlabeled because of the tracer position in glutamine (Fig. 4a, 4b).

Finally, this analysis gives an idea about how pyruvate is exported from the matrix. The appearance of additional intramatrix pyruvate increases the OA pool. As a result, the amount of citrate and α-KG is also increased. Additional citrate is exported more actively and leads to an increased level of OA in the cytosol; this additional OA may be partly converted into pyruvate via nonspecific decarboxylation, PEPCK-dependent decarboxylation, or through malate by cytosolic malic enzyme (ME1). Since ME1 knockdown did not affect the pyruvate export flux induced by Grubraw, the former pathways seem to be more probable.

We next tested whether the excessive mitochondrial pyruvate export was an effect characteristic of HeLa Kyoto cells only, or if it was typical for different cancer cell lines of the glycolytic metabolic type. We obtained cancer cell lines HT29 (human colorectal cancer), MDA-MB 231 (human triple-negative breast adenocarcinoma), WM164 and Lu451 (human melanoma), and non-cancer line H9C2 (immortalized rat cardiomyoblasts) stably expressing active and inactive Grubraw (Extended Data Fig. 5a – d). Under the same experimental conditions as HeLa cells, all cancer cell lines demonstrated Grubraw-dependent pyruvate export (Fig. 5a). On the contrary, in the H9C2 cell line the Grubraw-induced pyruvate export was far less pronounced. In melanoma cell lines, lactate export was proven to be independent of Grubraw activity (Fig. 5b). Moreover, Grubraw activity in melanoma cell lines led to an increase in OCR (Fig. 5c, 5d).

**Figure 5.**
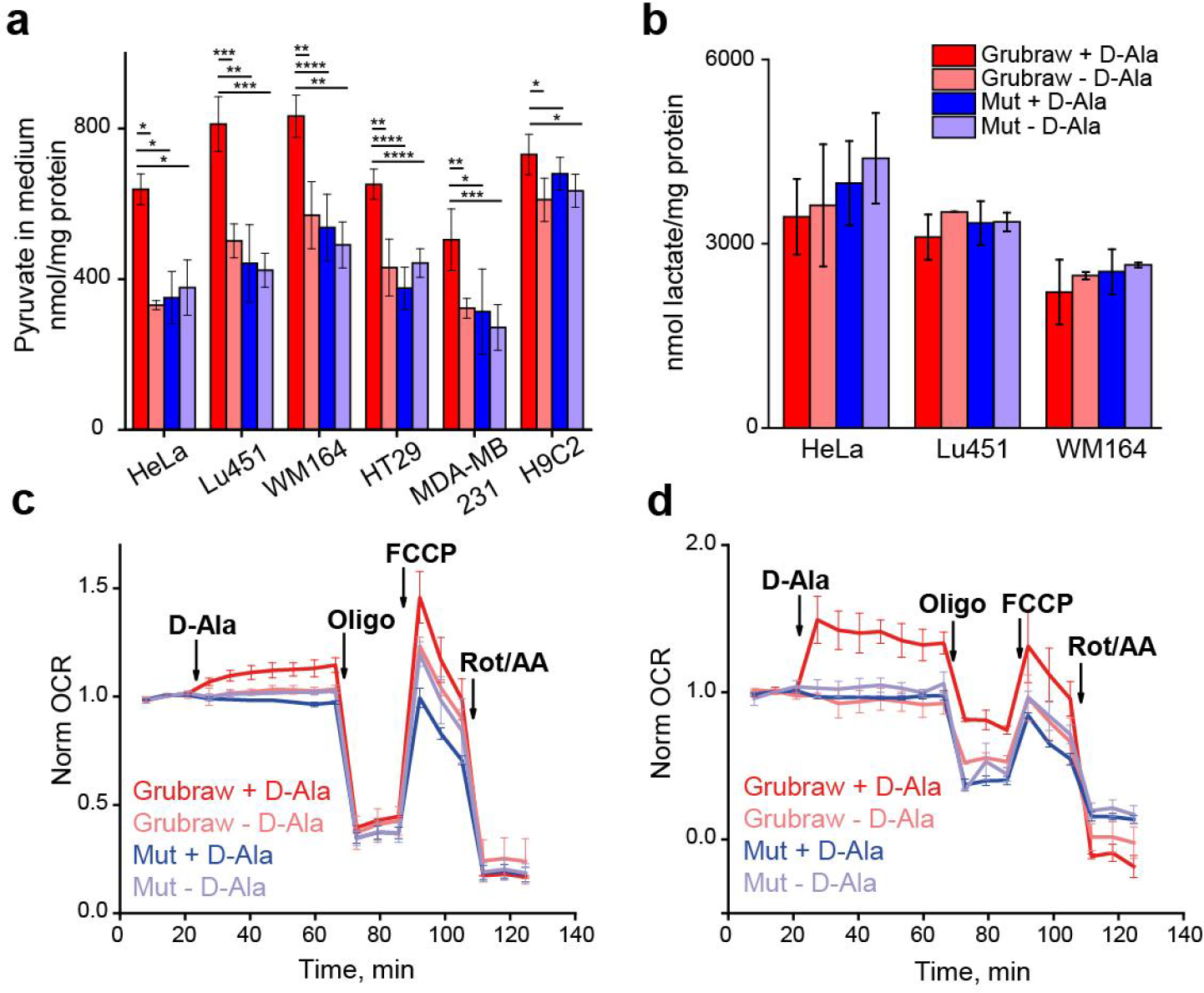
Grubraw activity modulates metabolism in different cancer cell lines. **a**, The effect of Grubraw activity on extracellular pyruvate in a panel of cancer cell cultures. For reference, the data for HeLa Kyoto cells from Fig. 2a are present. **b**, The effect of Grubraw activity on extracellular lactate in melanoma cultures. For reference, the data for HeLa Kyoto cells from Fig. 2c are present. **c**, The effect of Grubraw activity on OCR in Lu451 cells. **d**, The effect of Grubraw activity on OCR in WM164 cells. Data in **a** and b represent mean pyruvate or lactate quantity in the medium normalized to protein quantity in the sample ±SD. *p<0.05, **p<0.005,***p<0.0005, ****p<0.00005 In **c** and **d**, mean OCR normalized to the corresponding OCR in the zero time point ±SD is shown at every time point for 2-3 biological replicates for every sample.

Summarizing the data presented above, we have identified pyruvate release by cells as a specific mechanism associated with control of the level of pyruvate (or, more generally, control of the carbon flux) in the mitochondrial matrix of cells with Warburg-type metabolism. In oxidative metabolic type cells, like H9C2^22^, this mechanism functions much more weakly. We believe that this is a previously undescribed mechanism that allows cancer cells to control the activity of mitochondria in addition to the commonly accepted restriction of pyruvate intake to the matrix.

We next tested whether Grubraw activity affected tumor growth rate in the Lu451 xenograft model in immunodeficient Nude mice. Lu451 cells with stable firefly luciferase and active or mutant Grubraw expression were obtained using CRISPR genome editing (Extended Data Fig. 6a). Luciferase activity and Grubraw-dependent pyruvate export were proven in cell culture (Extended Data Fig. 6b, 6c). An *in vitro* proliferation test with or without D-Ala revealed a difference in cell division rate between Lu451-Fluc-P2A-Grubraw and the inactive mutant (Fig. 6a).

**Figure 6.**
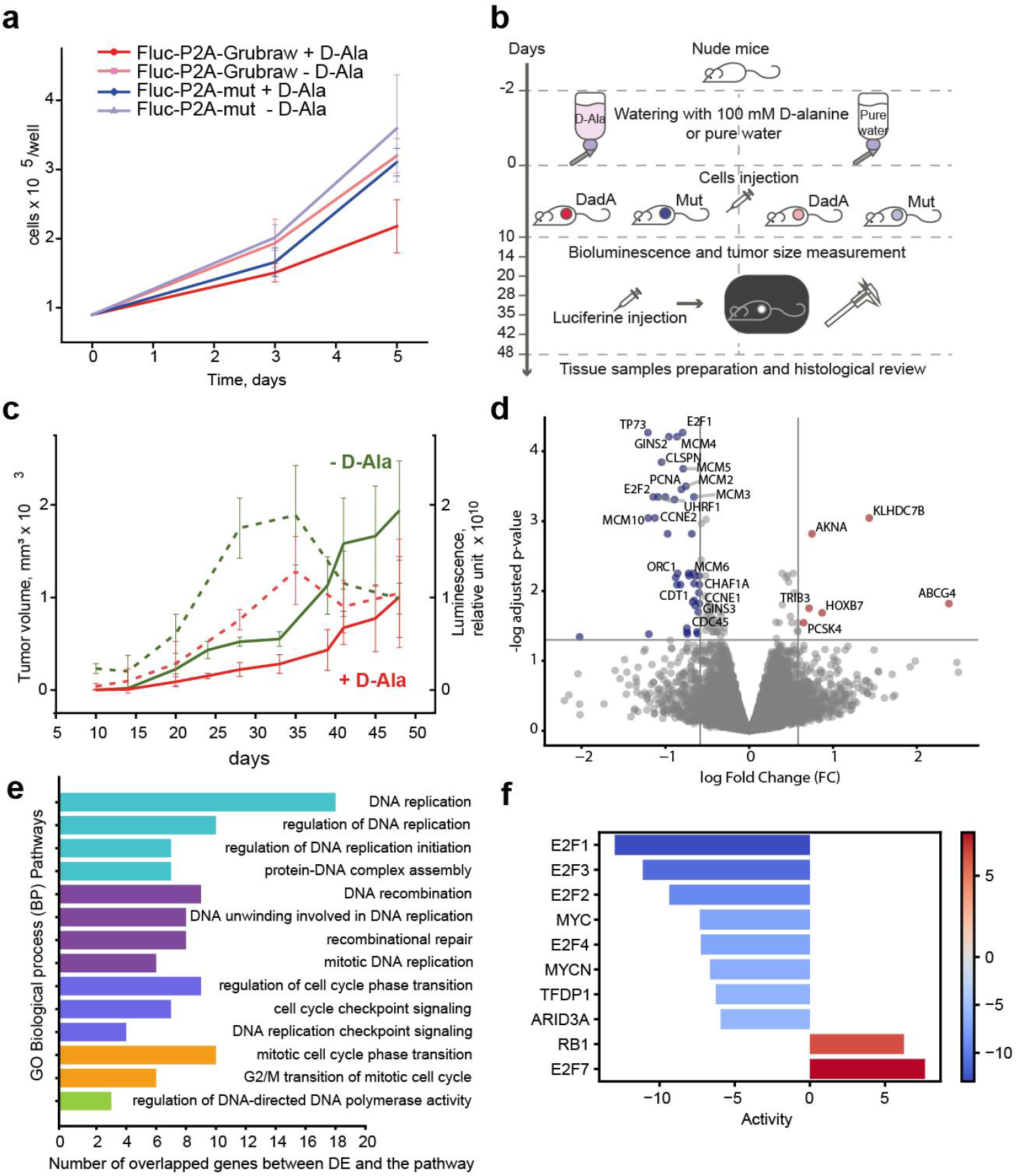
Grubraw activity slows down tumor growth *in vivo*. **a**, The effect of mRuby-Grubraw activity on Lu451 cell culture growth. **b**, The design for the experiment with the Lu451 xenograft model. **c**, The effect of Grubraw activity on tumor volume (solid lines) and bioluminescence (dashed lines). Mean values ±SD are shown. **d**, Volcano plot displays differentially expressed genes, up (red) and down (blue) upon activation of mitochondria. **e**, Results of gene set enrichment analysis for DE downregulated genes in Gene Ontology Biological process pathways. Color represents the corresponding parent source for each of the pathways in Gene Ontology: cyan - DNA-templated complex assembly initiation; violet - unwinding repair change recombination; blue - negative integrity checkpoint signaling; orange - G2/M mitotic transition phase; green - positive DNA-directed polymerase activity. **f**, Top 10 most modified transcription factors (TF) depicted from transcriptome profile upon activation of mitochondria by Grubraw.

The scheme of the experiment with mice is shown in Fig 6b. Depending on the group, mice with subcutaneously grafted cells received either 0.1M D-Ala in drinking water, or just pure water. In the control group with xenografts expressing mutant Grubraw, D-alanine did not affect tumor growth rate (Extended Data Fig. 6d). In the groups with xenografts expressing active Grubraw, D-Ala decreased tumor growth rate, especially 14 – 28 days after tumor cells injection. By day 28 the mean tumor volume in the group watered with D-Ala was about 2-fold smaller than in the group watered with pure water (Fig. 6c, Extended Data Fig. 6e). Tumor necrotization started 28 days after injection in the water drinking group and after 35 days in the group watered with D-Ala (Fig. 6c). Histological review of tissue sections 48 days after xenograft implantation did not reveal any microscopic changes in the lungs, liver, brain, nor lymph nodes of the experimental animals (Extended Data Fig. 6 g – k). On day 48, the size of necrosis reached up to 50% of the tumor size in the water group, and up to 30% in the D-alanine group (Extended Data Fig. 6k). After necrosis had started, the luminescence began to decrease in all groups, and the dispersion in tumor size inside the groups increased. No toxic effects of 0.1 M D-alanine, including significant weight loss (Extended Data Fig. 5f), were observed in these mice. Our results show that activated mitochondrial metabolism markedly slows down melanoma growth *in vivo*.

In order to understand the possible mechanisms that lead to decreased proliferation upon activation of mitochondria by Grubraw, we used RNA sequencing to investigate transcriptomic profiling shifts during Grubraw activation. We analyzed the same cell lines that were used in the experiments with xenografts. Cells expressing non-activated Grubraw or its inactive version had the same transcriptomic profile, and the addition of D-Ala to cells expressing inactive Grubraw also had no effect (Extended Data Fig. 7a).

In contrast, in the samples with activated Grubraw, differential expression analysis revealed significant downregulation of a narrow set of 41 genes that included key cell cycle regulators E2F1, E2F2, Cyclin E (both CCNE1, CCNE2), the core genes of CDC45-MCM-GINS (CMG) helicase complex (GINS2, GINS3, CDC45, MCM2-6, MCM10) and genes that regulate the initiation of replication: Origin Recognition Complex Subunit 1 (ORC1); Proliferating Cell Nuclear Antigen (PCNA); Chromatin Licensing And DNA Replication Factor 1 (CDT1) that is involved in the pre-replication complex; and Chromatin Assembly Factor 1 Subunit A (CHAF1A) (Fig. 6d).

Out of 41 downregulated genes, 25 were found in gene-enrichment analysis as significantly enriched in replication processes (Fig. 6e, Extended Data Table 3). In addition, we estimated which transcription factor-related pathways were changed by Grubraw activity and found that the five most significantly decreased pathways were related to E2F1, E2F2, E2F3, E2F4 and MYC. It was previously described that E2F genes contribute to the Warburg effect, restrict glucose oxidation and suppress mitochondrial biogenesis in proliferating cells^23^. Here we demonstrate that suppression in the opposite direction also works and mitochondrial activity downregulates E2F genes.

The pathways RB1 and E2F7, the well-known suppressors of cell cycle and proliferation, were increased by activity of Grubraw (Fig. 6f). We also observed downregulation of the activity of the kinase pathways MAPK, PI3K, and JAK-STAT, with parallel activation of the p53 pathway (Extended data Fig. 7b).

Among six upregulated genes were KLHDC7B and AKNA. The former was previously described as an endoplasmic reticulum stress activator leading to apoptosis^24^. The latter, AKNA, is an AT-hook transcription factor which regulates microtubule organization and cell delamination and hence influences the epithelial-to-mesenchymal transition^25^ (Fig. 6d).

Importantly, we also identified the Lactate Dehydrogenase A (LDHA) as a downregulated gene. It was shown before that MYC activated LDHA expression leading to glycolytic switch and tumor growth ^26^. Our experiments demonstrate that backward regulation also exists and the activation of mitochondria leads to both LDHA and MYC downregulation, together with suppression of key proliferation regulators.

In summary, we created a chemogenetic tool named **Grubraw** capable of activating mitochondrial metabolism. Mitochondrial activation with Grubraw involves the production of pyruvate and FADH_2_ by enzymatic oxidation of D-Ala in the matrix, increasing oxygen uptake following activation of the TCA cycle and ETC. Grubraw can be used to activate mitochondria of cells with low mitochondrial pyruvate intake, which is characteristic of various cancer cells^27–29^. Apparently, the activity of Grubraw associated with the transfer of electrons from FADH_2_ to quinones can partially compensate for disturbances in the operation of the electron transport chain associated with mutations or other pathological disorders of complex I.

We observed that the mitochondrial pyruvate produced by Grubraw is partially secreted into the extracellular medium by cancer cells, while in cardiomyoblasts with a more pronounced oxidative metabolism, this effect was very weak. This demonstrates that cells with Warburg-type metabolism are able to control mitochondrial carbon load not only by restricting pyruvate influx into the matrix, but also by actively exporting pyruvate produced in the mitochondria out of the matrix and the cell. We hypothesize that in the tumor microenvironment excessive carbon efflux from the mitochondria and cells not only restricts respiration, but also provides an additional carbon source to the other cells in the niche.

Another important observation is that activation of mitochondria in cancer cells does not induce oxidative stress. One of the frequently spoken theories suggests that cancer cells switch off mitochondria to diminish ROS formation. Our data question this explanation.

We found that reversing the Warburg effect by activating mitochondrial metabolism leads to a decrease in cancer cell proliferation and tumor growth rate. In order to explore possible mechanisms of this effect, we performed a transcriptomic analysis and demonstrated downregulation of key factors responsible for proliferation. It is well known that activation of proliferation in cancer cells driven by a variety of transcription factors, kinase cascades and metabolic changes suppresses mitochondrial metabolism. Our results are the first to demonstrate that the opposite direction of regulation is also possible. This gives an unexpected turn to possible explanations of the Warburg effect. Most of the existing explanations of the Warburg switch are “cancerocentric”. They try to explain why and how rapidly proliferating cancer cells benefit from decreasing mitochondrial respiration and metabolism, either by ensuring better biosynthesis, or by decreasing ROS formation, etc. Our data demonstrate that while cancer cells can downregulate mitochondrial functions, mitochondria can also downregulate cancer cell proliferation. This suggests a new explanation of the Warburg switch. Early in the evolution of eukaryotes, mitochondria were prokaryotic invaders, likely intracellular parasites in the beginning, with the following transition to symbiosis. When a parasite invades a cell, it reroutes cellular machinery to ensure optimal survival and multiplication within the host, at the expense of the host’s own functioning. Moreover, sometimes it is a strategy of the host to switch off some signaling/metabolic pathways when the activity of the parasite is detected. In addition, a number of defense strategies exist that decrease the activity of the parasite when the host has to function, e.g. replicate. Further in transition from parasitism to symbiosis these mechanisms could still exist to ensure a dynamic balance between the host’s and the symbiont’s needs. And even now, in modern eukaryotic cells, this mutual regulation between the “parasite/symbiont” (mitochondria) and the host (cell) may still exist. If the latter is true, in order to proliferate, eukaryotic cells should suppress the “parasite” or “symbiont” by, for example, restricting the substrate availability (pyruvate). And when the “parasite/symbiont” is active, proliferation of the “host” is suppressed.

Our hypothesis might explain why cancer cells reduce mitochondrial activity, and, in the opposite direction, mitochondria reduce cancer cell proliferation. This suggests that the Warburg effect probably has nothing to do with metabolic advantages, but instead repeats an evolutionarily inherited mechanism of host-parasite interaction. In favor of this suggestion is the fact that Warburg metabolism is characteristic not just of cancer cells, but also of other cells that need to rapidly multiply, like activated T-cells ^30^, proliferating cells in the developing embryo^11,31^, and proliferating stem and progenitor cells in adult organisms ^32–34^. Further studies are necessary to verify this hypothesis, but the results of this verification could bring new therapeutic strategies for tumor suppression.

## Acknowledgements

This paper is dedicated to the memory of Vladimir Skulachev and Andrey Vinogradov, two great Russian scientists who contributed enormously to the field of mitochondrial research. We want to acknowledge Lubov I. Golubeva and Ekaterina S. Kovaleva from Joint-Stock Company “Ajinomoto Genetika Research Institute” for advice in planning and interpreting mass isotopomer measurements.

This research was funded by the Russian Science Foundation (RSF), grant number 23-75-30023.

CRISPR genome editing was funded by the Ministry of Science and Higher Education of the Russian Federation grant number 075-15-2019-1789 to the Center for Precision Genome Editing and Genetic Technologies for Biomedicine.

## Author Contributions

E.S.P. and D.Y.B. designed the research; carried out biochemical, cell and microscopic experiments; designed molecular cloning; analyzed data from imaging, in vitro and in vivo experiments and co-wrote the paper. A.V.I. carried out CRISPR technologies. A.A.M. carried out lentivirus production. D.A.K., A.E.K. designed and carried out proteomics, mass spectrometry and metabolomics analyses. A.M.N. and O.M.K. designed and performed bioinformatic analyses and co-wrote the paper. N.F.Z. and A.V.I. carried out oxygen consumption rate measurement. L.E.S. and M.V.S. performed experiments with mice. O.V.L. carried out extracellular lactate measurement. O.I.P. performed immunohistochemistry and histological review of mouse tissues. I.B. helped with melanoma culturing. G.G., O.G., E.S. performed RNA sequencing analysis. V.V.B. directed the research and co-wrote the paper.

## Author Information

Correspondence and requests for materials should be addressed to E.S.P., or V.V.B.

## Data and code availability

LC-MS/MS data on ^13^C inclusion into intra- and extracellular ketoacids were uploaded on Mendeley Data (https://doi.org/10.17632/ws58h3crzh.1). GC-MS data on ^13^C inclusion into lactate and other metabolites were uploaded on Mendeley Data (https://doi.org/10.17632/9d65p4z799.1). Proteomics data on shRNA knock-down of single genes are deposited on PRIDE (ID: PXD044346). Example code for processing LC-MS data for ^13^C inclusion analysis is deposited on GitHub (https://doi.org/10.5281/zenodo.8241945).

RNASeq data is available in SRA via project PRJNA1025975.

Any additional information required to reanalyze the data reported in this paper is available from the lead contact upon request.

## Methods

### Gene cloning and mutagenesis

We synthesized cDNA of the *Pseudomonas aeruginosa* DadA gene (GenBank: AUB04911.1) at Evrogen (https://evrogen.com/). The DadA was cloned into the pQE30 vector using BamHI/SacI restriction sites (see primer list). The cDNA of DadA was fused with mRuby or EGFP via a Gly-Ser-Gly linker containing the BamHI restriction site and subcloned into the pQE30 vector using the BglII/SacI sites in the primers and BamHI/SacI in the vector.

For expression in the eukaryotic cell mitochondrial matrix, cDNAs of the FP-fused DadA were subcloned into the second generation lentiviral vector pLCMV-Puro with an N-terminal tandem dimer of COX8 presequences using BamHI site in the vector and BglII in the flanking primers. Primers with nucleotide substitutions leading to Tyr17Ala and Tyr18Ala in the predicted FAD binding domain of the DadA were generated. The mutated DadA gene fragment lacking 18 end base pairs was amplified using dadA-Y17A/Y18A-Fw and DadA-SacI-Rv primers. Then the missing base pairs were added using dadA-BamHI-long-Fw primer which overlapped with the 5`-end of dadA-Y17A/Y18A-Fw primer.

Primer sequences are presented in Supplementary in Table 1.

All genetic construct sequences were verified by Sanger sequencing.

### Multiple Sequence Alignment

The amino acid sequence of DadA from *Pseudomonas aeruginosa* (GenBank: AUB04911.1) was aligned in BLAST against the SwissProt database. 9 sequences of 60 – 70% similarity were selected, and multiple global alignment in Clustal W was performed using default parameters. For visualization, a logo was created using WebLogo 3 ^35^.

### Protein purification

DadA variants were expressed in *E. coli* strain XL1 Blue using pQE30 plasmid with the His6 tag on the N-end of target protein. Bacterial Protein Extraction Reagent (Thermo Scientific) was used for bacteria lysis. Centrifuged lysate was purified by gravity-flow chromatography on TALON Metal Affinity Resin (Clontech) following the manufacturer’s protocol with 0.05% Triton X-100 added to all buffers. Fractions containing target protein were desalted and concentrated on Amicon Ultra 10K centrifugal filter units (Merck Millipore), and the final protein preparation was stored in PBS containing 0.05% Triton X-100 at 4°C.

### SDS-PAGE

The purity and concentration of DadA preparations were evaluated using SDS-PAGE in 12% separating gel with Coomassie staining. Protein concentration was evaluated using BSA standards (100 – 1000 ng/lane) loaded in the same gel. The standard curve was plotted in ImageJ.

### DadA activity assay

DadA activity was measured in microplates as described in ^8^ with minor alterations. The reaction mix contained 100 mM Tris-HCl (pH 8.0), 0.8 mM phenazine methosulfate, 0.8 mM nitroblue tetrazolium, 20 mM D-Ala, 0.2 mM FAD (optionally) and 0.5 – 1 μg protein. All DadA preparations demonstrated the same activity with and without additional FAD. The D_620_ increase was registered at 37°C. A Varioscan Lux (Thermo Scientific) plate spectrophotometer was used in this and all following plate assays. To prove that metabolic inhibitors do not affect DadA

activity, 100 μM rotenone,120 μM CPI-613, 10 μM UK-5099, or 0.5 μM AR-C155858 was added to the reaction mix. All activities were normalized to wild type DadA activity without inhibitors.

To test whether DadA could reduce NAD^+^ or NADP^+^, the reaction mix containing 100 mM Tris-HCl (pH 8.0), 2 or 10 mM NAD^+^ or NADP^+^, 20 mM D-Ala, and 0.5 – 1 ug protein was used. The D_340_ increase was registered at 37°C.

All measurements were performed in 2 – 3 technical replicates.

### Pyruvate assay

The cell cultures with stable mRuby-Grubraw or mRuby-Grubraw-mut expression were incubated in HBSS with 20 mM HEPES and 11 mM glucose (except as noted). To inhibit MPC or MCT 1 and 2, 10 μM UK-5099 or 0.5 μM AR-C155858, respectively, were added to the medium. After 40 min incubation at 37°C and 5% CO_2_, 10 mM D-alanine was added, and the cells were incubated for 1 h 15 min. Pyruvate concentration in the centrifuged extracellular medium samples (5 min 18000 g, 4°C) was measured in microplates using Pyruvate Assay kit (Abcam) or as described in ^36^.

The cells were lysed in RIPA buffer, centrifuged (5 min 18000 g, 4°C), and protein concentration in the supernatant was measured using the BCA Protein Assay kit (Sigma).

Total pyruvate content in the well normalized to total protein content (5 – 7 biological replicates) was compared by t-test with Holm adjustment. For knockdown HeLa samples, these values were additionally normalized to the corresponding control value (Grubraw-mut without D-alanine) and compared using the Kruskal-Wallis rank sum test and Dunn test with Holm adjustment.

### Lactate assay

The samples were collected as described in pyruvate assay protocol. Lactate concentration was measured using Lactate Assay kit II (Sigma) or Radiometer ABL800 FLEX device. Total lactate content in the well was normalized to total protein content as described in the pyruvate assay protocol. 2 biological replicates were analyzed.

### Cell culturing

HeLa Kyoto cells (gender: F) were cultured in RPMI 1640 (Sigma) containing 10% FBS v/v (Biosera), 2 mM L-glutamine (Paneco), 100 U/mL penicillin (Paneco) and 100 mg/mL streptomycin (Paneco) at 37 °C in a 5% CO2 atmosphere.

Lu451 (gender: M) and WM164 (gender: M) cells were maintained in DMEM medium (Paneco) containing 4.5 g/L glucose and 5% FBS, 2 mM L-glutamine (Paneco), 100 U/mL penicillin (Paneco) and 100 mg/mL streptomycin (Paneco) at 37 °C in a 5% CO2 atmosphere.

HT29 (gender: F), MDA-MB 231 (gender: F), HEK 293 TN (gender: F), H9C2 cells were cultivated in DMEM medium (Paneco) containing 4.5 g/L glucose and 10% v/v FBS, 2 mM L-glutamine (Paneco), 100 U/mL penicillin (Paneco) and 100 mg/mL streptomycin (Paneco) at 37 °C in a 5% CO2 atmosphere.

### shRNA Cloning

shRNA plasmids to knockdown PDH1a, ME1, PC and MCT4 expression by RNA interference were created based on pGreenPuro lentivector (System Biosciences, LLC). The shRNA template was made by annealing appropriate oligonucleotides to each other (Extended Data Fig. 3) and then cloned into the unique BamHI or EcoRI sites in the vector.

### Generating cell lines with stable protein or shRNA expression

The HeLa Kyoto, HT29, MDA-MB 231, and H9C2 cell lines stably expressing Grubraw variants or shRNAs were generated by lentiviral transduction followed by cell sorting.

Lu451 and WM164 cell lines stably expressing Grubraw variants were generated by CRISPR genome editing followed by cell sorting.

### Lentivirus production

HEK 293 TN cells were seeded at 5 × 104 cells/cm^2^ in 25 cm^2^ tissue culture flasks. After 22 hours, cells were transfected using linear 40 KDa polyethyleneimine (Polysciences) at a mass ratio of 1:3 (DNA:PEI), with the respective plasmids: 5 μg of vector genome; 2.5 μg pMDLg/pRRE; 2.5 μg of pRSV-REV; 2.5 μg of VSV-G envelope. Medium was replaced 18-22 hours after transfection. After another 48-hour period, the supernatant was harvested, clarified at 0.45 μm and stored at −80 °C. The resulting virus concentration was about 2*10^6 TU/ml.

### Lentiviral transduction

The cells were seeded on culture plastic so that they were transduced at 60% confluency. Polybrene was added to a concentration of 8 μg/ml. After 2 – 3 days, the transduced cells began to fluoresce and were collected by cell sorting.

### Cell sorting

Cells suspended in ice-cold HBSS with 20 mM HEPES and 11 mM glucose were sorted on a Sony MA9000 Sorter and collected into 0.5 ml of cultivation medium. mRuby fluorescence was excited by a 561 nm laser, EGFP or GFP fluorescence was excited by a 488 nm laser. For cells expressing both fluorescent proteins (green and red), manual fluorescence spillover compensation was used.

### CRISPR genome editing

The intron of the PPP1R12C gene at the AAVS1 locus of chromosome 19 was chosen as the insertion site. A bicistronic lentiCRISPRv2-Blast (https://www.addgene.org/83480/) plasmid carrying SpCas9 was used, into which the guide RNA sequence was cloned under the U6 promoter at the BsmBI restriction sites. Double strand DNA for gRNA cloning was made by annealing oligonucleotides gRNAgRNA-AAVS1-2-dir and gRNA-AAVS1-2rev to each other (Suppl. Table 1).

A donor plasmid carrying the insertable construct between the homology arms was created based on the AAV-CAGGS-EGFP plasmid (https://www.addgene.org/22212/). EGFP was replaced by MCS containing homologous arms obtained by annealing the oligonucleotides Ins-dir-NcoI and Ins-rev-MluI to each other (Supp. Tabl.1). Overlap extension PCR was used for connection of firefly luciferase with mRuby-Grubraw or mRuby-Grubraw-mut by P2A self-cleaving peptide, and then PCR-products were introduced into the MCS region digested with AgeI and KpnI using GeneArt™ Gibson Assembly HiFi Master Mix according to manufacturer protocol.

Two-component transfection with the Grubraw donor plasmid and SpCas9/gRNA plasmid was used to transiently express SpCas9 and gRNA followed by editing the melanoma Lu451 cell genome. Five days after transfection, cells were selected for knock-in with 1.5 μg/ml puromycin. After selection, cells were validated for mRuby signal using FACS (Fluorescent-Activated Cell Sorting) (Sony MA900).

### qPCR

RNA was extracted from cultured cells using the ExtractRNA kit (Evrogen) according to the manufacturer’s protocol. OneTube RT-PCR SYBR kit (Evrogen) was used for cDNA production by reverse transcription followed by qPCR. Primers for the human housekeeping gene TUBA1B, and the PDH1a, ME1, PC, and MCT4 genes are described in Extended Data Fig. 3. Relative changes in genes expression were analyzed with the 2^-ΔΔCt^ method^37^.

### Transfection

For HeLa Kyoto cells, FuGENE HD Transfection Reagent (Promega) was used. The cells were seeded on plastic or confocal dishes 2 days before the experiment and transfected 1 day before the experiment.

### Microscopy

Fluorescent imaging was performed using an Eclipse Ti2-E microscope (Nikon), the NIS-Elements program (Nikon) and a monochrome Prime BSI digital camera (Teledyne Photometrics), if not otherwise stated. Image analysis and editing was performed in ImageJ and Nikon Analysis programs. The digital data were analyzed and plotted in Origin 2015. For imaging parameters, see Supplementary Table 2.

To confirm the mitochondrial localization of DadA, cells expressing active or inactive mRuby-Grubraw and EGFP-Grubraw were stained with MitoTracker Green (ThermoFisher Scientific) or20 nM TMRM, respectively. Mitochondrial membrane potential was also evaluated by TMRM fluorescence.

To confirm the matrix localization of DadA, HeLa Kyoto cells coexpressing mRuby-Grubraw and HyPer7-Micos60 were imaged on a Nikon Ti2 confocal microscope with a Nikon A1 LFOV monochrome digital camera.

H_2_O_2_ generation was evaluated using HyPer7 ^12^ with a mitochondrial matrix localization signal. In this and all further time-lapse experiments, HeLa Kyoto cells expressing mRuby-Grubraw or mRuby-Grubraw-mut were used. At the end of the imaging session, 100 μM H_2_O_2_ was added to prove sensor functioning.

Matrix pH dynamic was evaluated using SypHer3s ^15^ with a matrix localization sequence. NAD^+^/NADH ratio dynamics was registered using SoNar ^16^. At the end of the imaging session, 2.5 mM sodium lactate was added as a positive control to prove sensor functioning.

Pyruvate concentration dynamics in the cells was evaluated using Pyrates ^11^. To model physiological glycolysis flux and cytosolic pyruvate concentration, 11 mM glucose was added to the imaging medium. To inhibit PDH, 120 μM CPI-613 was added. To activate DadA, 10 mM D-alanine was added. At the end of the imaging session, 30 mM pyruvate was added to prove sensor functioning.

Glucose concentration dynamics in the cells was evaluated using FLII^12^Pglu-700μδ6 ^17^. The cells were seeded into μ-slides (iBiDi). The culture medium was replaced with imaging medium with 10 mM D-alanine and 11 mM D-Glucose 30 min before the experiment. After the baseline sensor signal was registered, the medium was replaced by imaging medium with 10 mM D-alanine. Imaging parameters were the same as for the Pyrates sensor.

In all experiments, the cells were incubated in imaging medium (HBSS medium with 20 mM HEPES) for 30 min at 37°C and 5% CO_2_, then the baseline sensor signal was registered for ∼5 min, after which 10 mM D-alanine was added (if not otherwise stated). Sensor fluorescence ratio or TMRM fluorescence intensity in every cell was normalized to the corresponding mean value for the baseline in this cell. These normalized values were averaged for n cells. For HyPer7 and FLII12Pglu-700μδ6, the non-normalized sensor fluorescence ratio was averaged for n cells. All experiments were performed in 2 – 3 replicates.

Murine tumor and tissue sections were imaged in brightfield mode with a Nikon DS-Fi3 Color Camera and 20x objective.

### Oxygen consumption rate measurement

OCR was measured using Seahorse Mito Stress Test on Seahorse XF Analyzer (Agilent) according to the manufacturer’s protocol with minor alterations. Cells expressing mRuby-Grubraw or mRuby-Grubraw-mut were incubated in HBSS or cultivation medium without HCO_3_ ^-^ for 30 min at 37°C without CO_2_ . Basal OCR was registered for 18 min. Then 10 mM D-alanine or HBSS was added and OCR was registered for 42 min. Subsequently 1 μM oligomycin, 1 μM FCCP, and 1 μM antimycin + 1 μM rotenone were added, with 18 min OCR measurement after every administration, if not described else. OCR data for each well were normalized to basal OCR. Data for 3-5 biological replicates (wells) were averaged.

### LC-MS and LC-MS/MS analysis of ketoacids

Samples were derivatized with phenylhydrazine (PH) using a procedure adapted from ^38^. The derivatizing reagent (DR) was a freshly prepared 2 mM solution of PH base in acetonitrile-methanol-water mixture (2:2:1 v/v). For intracellular samples the cultural medium was removed and cells were incubated with 1 ml of DR (1h, -20°C). For extracellular samples an aliquot of cultural medium was mixed with DR (1:9) and incubated (1h, -20°C). Then the liquid phases were centrifuged and dried under vacuum (25°C) and dissolved in solvent A. The column in the HPLC system was a Thermo C18 Hypersil Gold 100×2.1mm, 1.9 μm. SolventA consisted of 5% acetonitrile, 0.3% triethylammonium formate (TEA) in water, and solvent B consisted of 0.3% TEA in acetonitrile. The following gradient was used: 0-3 min: isocratic 5%B, 23 min: 45%B, 24-29 min: isocratic 90% B. The mass-spectrometer (Thermo Q-Exactive HF, HESI-II ion source) was operated in negative ion mode. To register MS1 for PH derivatives we used single ion monitoring mode with 60 000 resolution at three m/z ranges: 176.066-180.066 (Pyr); 220.100-224.100 (OA); 234.100-238.100 (α-KG). MS2 spectra of these substances were registered in separate runs of parallel reaction monitoring mode for precursor ions: 177.066, 178.070, 179.073, 221.056, 222.060, 223.063, 235.072, 236.075, 237.0786, 238.0795 m/z at 30,000 resolution and HCD energy 10 (Extended Data Fig. 4b). For absolute quantitation of extracellular pyruvate and α-KG, we prepared a series of pure substance dilutions in a fresh cultural medium. MIDs were extracted from MS1 runs and corrected to natural isotope occurrence utilizing MS2 data. Additionally, we extracted the probabilities of tracer inclusion into the first position in a molecule from MS2 data . Details on spectra processing, the equation for MID correction, and statistical inference are provided in the Supplementary Methods.

### GC-MS analysis

The cell medium was treated with cold methanol (4:1), refrigerated (1 hour, -22°C) and centrifuged (14000 g, 10 min). 1.5 ml of each supernatant were transferred to a new microcentrifuge tube and derivatized with MSTFA as described in ^39^. D-Norvaline was added before the procedure as an internal standard for concentration measurements. The GC–MS system was a Shimadzu TQ8040 GC–MS, with an Agilent CP-Sil 8 CB (50m × 0.25 mm i.d. × 0.25 μm) column. The carrier gas was UHP-grade helium at 0.88 mL/min. GC–MS analysis of the prepared samples was performed as described in ^39^. 0.5 μL was injected in the split mode at a ratio of 1:50 with two or three replicates. For control of the GC–MS system, data collection and initial preprocessing, the Shimadzu GCMS Postrun Analysis software was used. Mass isotopomer distributions (MIDs) for lactate were extracted from raw spectra (Extended Data Fig. 4d) using self-made scripts. Procedures and tools used for identification, peak integration and MID transformations are described in the Supplementary Methods.

### Targeted proteomics

Cells were treated with cold 80% methanol (30 min), scraped and centrifuged (10000 g, 5 min). The pellet was washed with cold acetone, resuspended in 100 mM (NH_4_)HCO_3_ (pH 8.0, 0.2% w/v Promega ProteaseMax) and sequentially treated with TCEP (5 mM, 1h, 50^o^C), iodoacetamide (15 mM, 1h, RT) and Trypsin Gold (1 μg, 3h, 37^o^C). Then, formic acid was added (up to 1% v/v), the samples were centrifuged (10000 g, 5 min), supernatants transferred to new tubes and vacuum-dried. Aliquots were taken from supernatants for peptide measurement and also dried. Dried samples were dissolved in the solution (2% acetonitrile, 0.05% TFA). The mass-spectrometer was a Thermo Q-Exactive HF-X. The HPLC system was a Thermo UltiMate 3000. The chromatography conditions used were the following Column - Thermo Acclaim PepMap RSLC 0.075×150 mm; Solvent A - 0.1% formic acid in water; Solvent B - 80% acetonitrile and 0.1% formic acid in water; Gradient - 4%-45% B in 90 min, 90-100 min - isocratic 95% B; Flow - 300 nanoliters/min. For mass spectra registration the data dependent mode was used and the MS1 range was 350-1500 *m/z*, the number of precursors was 20, and the MS2 range was 200-2000 *m/z*. Separate runs of the samples were performed under the same chromatographic conditions in single ion monitoring mode of several peptides originating from the proteins of interest. Raw data were processed using the standard proteomics protocol and then the peaks of most abundant peptides were quantified separately using self-made scripts (the details are described in the Supplementary Methods).

### Proliferation test

Hela Kyoto cell lines were seeded on cell culture dishes at a density of 8000 cells/cm^2^, and Lu451 lines at a density of 10000 cells/cm^2^. The concentration of D-glucose in the growth medium was 1 g/l. The number of cells in the wells was counted using Luna Cell Counter (ThermoFisher Scientific) with Trypan blue staining. 3-6 biological replicates were analyzed.

### Murine xenograft model

The experiments were carried out on 20 female, 6-week-old, athymic nude mice, purchased from the SPF animal facility of the Institute of Biology and Biomedicine of Lobachevsky State University of Nizhny Novgorod (Russia). All animal experiments were approved by the Ethics Committee of the Privolzhsky Research Medical University (Nizhny Novgorod, Russia), approval #6 from April 17, 2019.

Lu451 cells with CRISPR/Cas9 generated stable firefly luciferase and mRuby-Grubraw or mRuby-Grubraw-mut expression were used. 1.5×10^6^ cells in serum-free DMEM were injected subcutaneously into Nude mice. 2 groups of mice with mRuby-Grubraw or mRuby-Grubraw-mut expressing xenografts were watered with 0.1 M D-Ala from 2 days before the injection untill the end of the experiment (48 days after the injection), while 2 identical groups receivedpure water as control. Every group consisted of 4 – 5 animals. The mice were anesthetized with 2% isoflurane before tumor measurement. Xenograft tumor diameter was measured every 4—7 days with a caliper. Tumor bioluminescence was imaged using an IVIS Spectrum imaging system (Caliper Life Sciences, Hopkinton, MA, USA) every week from day 10 after cell implantation. Luciferin solution (150 mg/kg) was administered intraperitoneally 10 minutes before the visualization. The images were analyzed using Living Image software. After the end of the experiment, the mice were weighed, euthanized by an overdose of isoflurane, and the tumors, brains, lungs, and livers were isolated and fixed in buffered formaldehyde and delivered to histopathological expertise.

### Luciferase assay in cells

HBSS with 0.3 mg/ml luciferine was added to Lu451 cells with stable firefly luciferase and mRuby-Grubraw or mRuby-Grubraw-mut expression in a 96-well plate and confluence Bioluminescence kinetics was registered for 5 min.

### Immunohistochemistry of mouse tissue sections

The organs removed during necropsy (tumor with lymph nodes, liver, lung, brain) were placed in a buffered neutral 10% formalin solution for fixation for 24 hours. After standard histological processing based on isopropyl alcohol (10% buffered formalin - 2 hours, isopropyl alcohol - 18 hours, liquid paraffin - 3 hours), tissue fragments were placed into paraffin blocks. Histological sections with a thickness of 3-4 μm were made using rotary microtomes (CUT4062, SLEE medical, Germany), stained with hematoxylin and eosin according to the standard procedure (dewaxing: ortho-xylene, isopropyl alcohol, staining: Mayer hematoxylin - 8 minutes, watery eosin - 3 minutes, rinsing with running water, dehydration stage: isopropyl alcohol, ortho-xylene) and were enclosed under a cover film (Tissue-Tek Prisma histostainer, Tissue-Tek PrismaFilm, Sacura, Japan).

### RNA Sequencing and data processing

Lu451 cells with CRISPR/Cas9 generated stable firefly luciferase and mRuby-Grubraw or mRuby-Grubraw-mut expression were used. 0.5 × 10^6^ cells were seeded onto 60 mm diameter dishes and grown for 3 days with or without 10 mM D-alanine in DMEM with 1 g/L glucose, 5% FBS and no pyruvate. RNA was extracted according to the manufacturer’s instructions using the Qiagen RNeasy Mini Kit (Qiagen). The quality of total RNA was evaluated using a Bioanalyzer 2100 (Agilent). The quantity and purity of RNA was estimated on a NanoPhotometer (Implen). 800 ng of RIN ≥7 of total RNA was used for library construction using a NEBNext® Poly(A) mRNA Magnetic Isolation Module and NEBNext® Ultra II™ Directional RNA Library Prep Kit for Illumina (New England Biolabs) according to the manufacturer’s instructions. The quality of libraries was verified using the Bioanalyzer 2100 (Agilent) and the yield was validated by qPCR. Libraries were then sequenced on a NovaSeq6000 (Illumina) with pair-end 61 bp reading.

For bioinformatics analysis, reads were aligned on transcriptome reference from Gencode Release 43 (GRCh38.p13) by STAR ^40^ and quantified by FeatureCounts ^41^. Differential expression analysis was calculated using the limma R package with thresholds 1.5 for Fold Change and 0.05 for adjusted p-value. Enrichment analysis was conducted using the clusterProfiler R package ^42^. TF activity was calculated by decoupleR ^43^ and pathways were estimated by Progeny ^44^.

