## Supplementary Information for "Grubraw, a chemogenetic mitochondrial activator, reveals new mechanisms underlying the Warburg effect"

### Extended Data

**Table 1. Primer sequences**

| Primer name | Primer sequence (5’ - 3’) |
|---|---|
| DadA-SacI-Rv | TACCGAGCTCTTAGTGTGCTGGCGCTGGA |
| DadA-BamHI-Fw | ATTAGGATCCCGAGTTCTGGTCCTTGGCA |
| BglII-mRuby-Fw | AGTCAGATCTACCGGTCGCCACCGTGTCTAAGGGCGAAGAGCT |
| DadA-BglII-Rv | AGTCAGATCTTTAGTGTGCTGGCGCTGGATG |
| BglII-EGFP-Fw | AGTCAGATCTGTGAGCAAGGGCGAGGAG |
| BglII-EGFP-Fw | AGTCAGATCTACCGGTCGCCACCGTGAGCAAGGGCGAGGAG |
| dadA-Y17A/Y18A-Fw | AGCGGTGTCATCGGTACCGCCAGTGCGGCAGCACTGGCCCGTGCC |
| dadA-BamHI-long-Fw | TCACGGATCCCGAGTTCTGGTCCTTGGCAGCGGTGTCATC |
| gRNA-AAVS 1-2-dir | CACCGTGTC CCTAGTGGCCCCACTG |
| gRNA-AAVS 1-2rev | AAACCAGTGGGGCCACTA GGGACAC |
| Ins-dir-NcoI | CATGGAGATGACCGGTGTCTTAAGCTGGTACCCAATTGTTATCGATCGTGA |
| Ins-rev-MluI | CGCGTCACGATCGATAACAATTGGGTACCAGCTTAAGACACCGGTCATCTC |

**Table 2. Fluorescence imaging parameters**

| Parameter measured | Fluorophore | Excitation source | Fluorescence filters | Objective | Time-lapse (frame/min) |
|---|---|---|---|---|---|
| DadA localization (Nikon Eclipse Ti2-E) | MitoTracker Green, EGFP | 470 nm LED | Multi-band filter LED-DA/FI/TR/Cy5-4X-B-000 (Semrock) and emission filter 515/30 | 60x oil immersion objective and additional 2x lens | – |
| DadA localization (Nikon Eclipse Ti2-E) | TMRM, mRuby | 555 nm LED | Multi-band filter LED-DA/FI/TR/Cy5-4X-B-000 (Semrock) and emission filter 595/31 nm | 60x oil immersion objective and additional 2x lens | – |
| DadA localization (Nikon Ti2 confocal microscope) | mRuby | 561 nm laser | 561 nm dichroic mirror and 595/50 emission filter | 100x oil immersion objective | – |
| DadA localization (Nikon Ti2 confocal microscope) | HyPer7-Micos60 (pCS2+-HyPer7-Micos60 plasmid transfection) | 488 nm laser | 488 nm dichroic mirror and 525/50 emission filter | 100x oil immersion objective | – |
| Mitochondrial membrane potential | TMRM | 555 nm LED | Multi-band filter LED-DA/FI/TR/Cy5-4X-B-000 (Semrock) and emission filter 595/31 nm | 40x | 2 |
| $H_2O_2$ generation | HyPer7 (pCS2+-HyPer7-MLS plasmid transfection) | 395 and 470 nm LED | Multi-band filter LED-DA/FI/TR/Cy5-4X-B-000 (Semrock) and emission filter 515/30 | 40x | 2 |

| | | | | | |
|---|---|---|---|---|---|
| pH | SypHer3s (pC1-CMV-tdmito-SypHer3s plasmid transfection) | 395 and 470 nm LED | Multi-band filter LED-DA/FI/TR/Cy5-4X-B-000 (Semrock) and emission filter 515/30 | 40x | 2 |
| NAD+/NADH ratio | SoNar (pCS2+-SoNar plasmid transfection) | 395 and 470 nm LED | Multi-band filter LED-DA/FI/TR/Cy5-4X-B-000 (Semrock) and emission filter 515/30 | 40x | 2 |
| Pyruvate concentration | Pyrates (pCKI-Pyrates or pCKI-Pyrates-mito plasmid transfection) | 440 nm LED | Multi-band filter LED-CFP/YFP/mCherry-3X-A-000 (Semrock), emission filters 474/27 nm and 544/24 nm | 40x | 2 |
| Glucose concentration | pFLII12Pglu-700μ Δ6 (pFLII12Pglu-700μ Δ6 plasmid transfection) | 440 nm LED | Multi-band filter LED-CFP/YFP/mCherry-3X-A-000 (Semrock), emission filters 474/27 nm and 544/24 nm | 40x | 2 |

**Table 3. Enriched Gene Ontology pathways, Biological Processes decreased under Grubraw activation (Extended data Table 3)**

| ID | Description | Gene Ratio | BgRatio | pvalue | p.adjust | qvalue | List of genes | Count | Category |
|---|---|---|---|---|---|---|---|---|---|
| GO:0006260 | DNA replication | 18/41 | 285/17133 | 2.59E-20 | 7.59E-18 | 5.77E-18 | MCM4/GINS2/MCM5/MCM2/PCNA/MCM3/MCM10/CCNE2/DTL/MCM6/ORC1/CHAF1A/CCNE1/GMNN/CDT1/GINS3/RMI2/CDC45 | 18 | DNA-templated complex assembly initiation |
| GO:0006261 | DNA-templated DNA replication | 15/41 | 155/17133 | 9.43E-20 | 1.84E-17 | 1.40E-17 | MCM4/GINS2/MCM5/MCM2/PCNA/MCM3/MCM10/CCNE2/MCM6/ORC1/CCNE1/GMNN/CDT1/GINS3/CDC45 | 15 | DNA-templated complex assembly initiation |
| GO:0006270 | DNA replication initiation | 13/41 | 37/17133 | 3.75E-25 | 2.19E-22 | 1.67E-22 | MCM4/MCM5/MCM2/MCM3/MCM10/CCNE2/MCM6/ORC1/CCNE1/GMNN/CDT1/GINS3/CDC45 | 13 | DNA-templated complex assembly initiation |
| GO:0090329 | regulation of DNA-templated DNA replication | 10/41 | 53/17133 | 1.95E-16 | 2.85E-14 | 2.17E-14 | MCM4/GINS2/MCM5/MCM2/PCNA/MCM3/MCM6/GMNN/CDT1/GINS3 | 10 | DNA-templated complex assembly initiation |
| GO:0006275 | regulation of DNA replication | 10/41 | 147/17133 | 7.86E-12 | 4.18E-10 | 3.18E-10 | MCM4/GINS2/MCM5/MCM2/PCNA/MCM3/MCM6/GMNN/CDT1/GINS3 | 10 | DNA-templated complex assembly initiation |
| GO:0033260 | nuclear DNA replication | 9/41 | 37/17133 | 5.63E-16 | 5.33E-14 | 4.06E-14 | MCM4/MCM2/PCNA/MCM3/MCM6/GMNN/CDT1/GINS3/CDC45 | 9 | DNA-templated complex assembly initiation |
| GO:0044786 | cell cycle DNA replication | 9/41 | 40/17133 | 1.23E-15 | 9.00E-14 | 6.85E-14 | MCM4/MCM2/PCNA/MCM3/MCM6/GMNN/CDT1/GINS3/CDC45 | 9 | DNA-templated complex assembly initiation |
| GO:0030174 | regulation of DNA-templated DNA replication initiation | 7/41 | 14/17133 | 2.90E-15 | 1.88E-13 | 1.43E-13 | MCM4/MCM5/MCM2/MCM3/MCM6/GMNN/CDT1 | 7 | DNA-templated complex assembly initiation |
| GO:0065004 | protein-DNA complex assembly | 7/41 | 225/17133 | 2.86E-06 | 5.97E-05 | 4.55E-05 | MCM2/CHAF1A/ASF1B/ASF1B/GMNN/CDT1/CDC45 | 7 | DNA-templated complex assembly initiation |
| GO:0000086 | G2/M transition of mitotic cell cycle | 6/41 | 140/17133 | 2.47E-06 | 5.36E-05 | 4.08E-05 | CLSPN/DTL/WEE1/ORC1/CDC25A/PKMYT1 | 6 | G2/M mitotic transition phase |
| GO:0044772 | mitotic cell cycle phase transition | 10/41 | 442/17133 | 3.21E-07 | 9.38E-06 | 7.14E-06 | E2F1/CLSPN/CCNE2/DTL/WEE1/ORC1/CDC25A/CCNE1/CDT1/PKMYT1 | 10 | G2/M mitotic transition phase |
| GO:0045786 | negative regulation of cell cycle | 9/41 | 394/17133 | 1.25E-06 | 3.17E-05 | 2.41E-05 | E2F1/CLSPN/WDR76/DTL/WEE1/ORC1/GMNN/CDT1/CDC45 | 9 | negative integrity checkpoint signaling |
| GO:1901987 | regulation of cell cycle phase transition | 9/41 | 425/17133 | 2.32E-06 | 5.23E-05 | 3.98E-05 | E2F1/CLSPN/WDR76/DTL/WEE1/ORC1/CDC25A/CDT1/CDC45 | 9 | negative integrity checkpoint signaling |
| GO:1901988 | negative regulation of cell cycle phase | 8/41 | 253/17133 | 4.59E-07 | 1.28E-05 | 9.72E-06 | E2F1/CLSPN/WDR76/DTL/WEE1/ORC1/CDT1/CDC45 | 8 | negative integrity checkpoint signaling |

| | | | | | | | | | |
|---|---|---|---|---|---|---|---|---|---|
| | transition | | | | | | | | |
| GO:0010948 | negative regulation of cell cycle process | 8/41 | 303/17133 | 1.78E-06 | 4.17E-05 | 3.17E-05 | E2F1/CLSPN/WDR76/DTL/WEE1/ORC1/CDT1/CDC45 | 8 | negative integrity checkpoint signaling |
| GO:0031570 | DNA integrity checkpoint signaling | 7/41 | 137/17133 | 1.00E-07 | 3.26E-06 | 2.48E-06 | E2F1/CLSPN/WDR76/DTL/ORC1/CDT1/CDC45 | 7 | negative integrity checkpoint signaling |
| GO:0000075 | cell cycle checkpoint signaling | 7/41 | 190/17133 | 9.26E-07 | 2.46E-05 | 1.87E-05 | E2F1/CLSPN/WDR76/DTL/ORC1/CDT1/CDC45 | 7 | negative integrity checkpoint signaling |
| GO:0000076 | DNA replication checkpoint signaling | 4/41 | 16/17133 | 9.63E-08 | 3.26E-06 | 2.48E-06 | CLSPN/ORC1/CDT1/CDC45 | 4 | negative integrity checkpoint signaling |
| GO:1900262 | regulation of DNA-directed DNA polymerase activity | 3/41 | 11/17133 | 3.35E-06 | 6.54E-05 | 4.98E-05 | GINS2/PCNA/GINS3 | 3 | positive DNA-directed polymerase activity |
| GO:1900264 | positive regulation of DNA-directed DNA polymerase activity | 3/41 | 11/17133 | 3.35E-06 | 6.54E-05 | 4.98E-05 | GINS2/PCNA/GINS3 | 3 | positive DNA-directed polymerase activity |
| GO:0006310 | DNA recombination | 9/41 | 327/17133 | 2.62E-07 | 8.08E-06 | 6.15E-06 | MCM4/GINS2/MCM5/MCM2/MCM3/MCM6/UNG/RMI2/CDC45 | 9 | unwinding repair change recombination |
| GO:0006268 | DNA unwinding involved in DNA replication | 8/41 | 22/17133 | 6.38E-16 | 5.33E-14 | 4.06E-14 | MCM4/GINS2/MCM5/MCM2/MCM3/MCM6/GINS3/CDC45 | 8 | unwinding repair change recombination |
| GO:0032508 | DNA duplex unwinding | 8/41 | 90/17133 | 1.34E-10 | 6.54E-09 | 4.98E-09 | MCM4/GINS2/MCM5/MCM2/MCM3/MCM6/GINS3/CDC45 | 8 | unwinding repair change recombination |
| GO:0032392 | DNA geometric change | 8/41 | 96/17133 | 2.27E-10 | 1.02E-08 | 7.76E-09 | MCM4/GINS2/MCM5/MCM2/MCM3/MCM6/GINS3/CDC45 | 8 | unwinding repair change recombination |
| GO:0071103 | DNA conformation change | 8/41 | 104/17133 | 4.33E-10 | 1.81E-08 | 1.38E-08 | MCM4/GINS2/MCM5/MCM2/MCM3/MCM6/GINS3/CDC45 | 8 | unwinding repair change recombination |
| GO:0000724 | double-strand break repair via homologous recombination | 8/41 | 161/17133 | 1.40E-08 | 5.46E-07 | 4.16E-07 | MCM4/GINS2/MCM5/MCM2/MCM3/MCM6/RMI2/CDC45 | 8 | unwinding repair change recombination |
| GO:0000725 | recombinational repair | 8/41 | 165/17133 | 1.70E-08 | 6.21E-07 | 4.72E-07 | MCM4/GINS2/MCM5/MCM2/MCM3/MCM6/RMI2/CDC45 | 8 | unwinding repair change recombination |
| GO:0006302 | double-strand break repair | 8/41 | 292/17133 | 1.35E-06 | 3.29E-05 | 2.51E-05 | MCM4/GINS2/MCM5/MCM2/MCM3/MCM6/RMI2/CDC45 | 8 | unwinding repair change recombination |

| | | | | | | | | | |
|---|---|---|---|---|---|---|---|---|---|
| GO:0000727 | double-strand break repair via break-induced replication | 7/41 | 11/17133 | 2.81E-16 | 3.28E-14 | 2.50E-14 | MCM4/GINS2/MCM5/MCM2/MCM3/MCM6/CDC45 | 7 | unwinding repair change recombination |
| GO:1902969 | mitotic DNA replication | 6/41 | 14/17133 | 1.03E-12 | 6.04E-11 | 4.60E-11 | MCM4/MCM2/MCM3/MCM6/GINS3/CDC45 | 6 | unwinding repair change recombination |

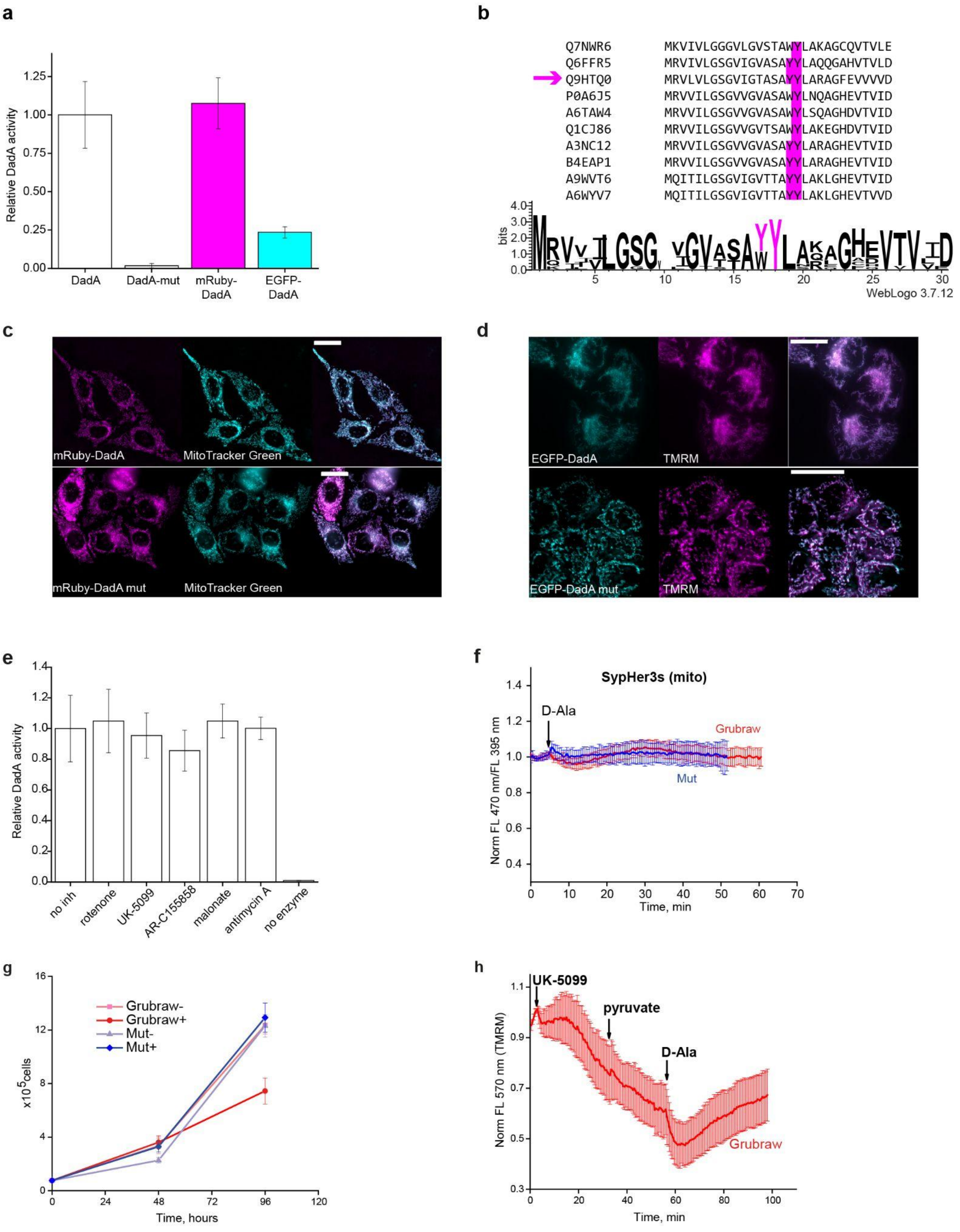


**Extended Data Figure 1. Metabolic instrument design. a,** Relative activities of wild-type DadA, DadA-mut (Y17A/Y18A point mutation), mRuby-DadA fusion, and EGFP-DadA fusion. Mean activities normalized to wild-type DadA activity ±SD are shown. **b,** Distribution of the amino acid residues in the putative FAD-binding N-end domain of DadA family proteins. For the sequences used in the multiple alignment, Uniprot identifiers are shown. DadA from the *P. aeruginosa* sequence is marked with a magenta arrow. The conservative tyrosine residues are shown in magenta. **c,** A fluorescence image of HeLa Kyoto cells expressing mito-mRuby-DadA or mito-mRuby-DadA-mut stained with mitoTracker Green. **d,** A fluorescence image of HeLa Kyoto cells expressing mito-EGFP-DadA or mito-EGFP-DadA-mut stained with TMRM. **e**, Relative enzymatic activity of wild-type DadA in presence of 100 μM rotenone, 10 μM UK-5099, 0.5 μM AR-C155858, 5 mM malonate, and 10 μM antimycin A . Mean activities normalized to control (DadA activity without inhibitors) ±SD are shown. All measurements were performed in 2 – 3 technical replicates. **f,** Mitochondrial matrix pH by Sypher3s sensor localized in the mitochondria. n = 18 – 20. **g,** HeLa Kyoto culture proliferation rate. Mean cell numbers in the wells for 3 biological replicates ±SD are shown. **h,** Evaluation of mitochondrial membrane potential in presence of 10 μM UK-5099 by TMRM for HeLa Kyoto cells expressing Grubraw. TMRM fluorescence intensity in every cell was normalized to the mean baseline value in the cell and averaged for 5 cells. In **c** and **d**, scale bars represent 25 μm.

**a. RNA interference and relative mRNA expression**

| GOI | Relative mRNA expression (normalized GOI amount relative to scramble control $2^{-\Delta\Delta Ct}$) | shRNA oligonucleotides (5′ - 3′) |
|---|---|---|
| PDH1a | 0,03 (0,01-0,06) | Fw: gatccGCCTGTAAGTATAATGGAATTCAAGAGATTCCATTATACTTACAGGCTTTTTTg<br>Rv: aattcAAAAAAGCCTGTAAGTATAATGGAATCTCTTGAATTCCATTATACTTACAGGCg |
| ME1 | 0,2 (0,17 - 0,25) | Fw: gatccGGAAGCCAAGAGGTCTCTTTATTCAAGAGATAAAGAGACCTCTTGGCTTCCTTTTTTg<br>Rv: aattcAAAAAAGGAAGCCAAGAGGTCTCTTTATCTCTTGAATAAAGAGACCTCTTGGCTTCCg |
| MCT4 | 0,11 (0,07 - 0,16) | Fw: gatccGAGCATCATCCAGGTCTACTTCAAGAGAGTAGACCTGGATGATGCTCTTTTTTg<br>Rv: aattcAAAAAAGAGCATCATCCAGGTCTACTCTCTTGAAGTAGACCTGGATGATGCTCg |
| PC | 0,23 (0,2 - 0,26) | Fw: gatccGTGAGATTGCCATCCGTGTTTCAAGAGAACACGGATGGCAATCTCACTTTTTTg<br>Rv: aattcAAAAAAGTGAGATTGCCATCCGTGTTCTCTTGAAACACGGATGGCAATCTCACg |

| GOI | qPCR primers (5′ - 3′) |
|---|---|
| PDH1a | Fw: TGCTGCTAACCAGGGCCA<br>Rv: AGACGTTCCCATTCCATAGCG |
| ME1 | Fw: GGCCTTTACCCTGGAAGAGAG<br>Rv: ATCCATTAAGAGAAGATACCTGTCA |
| MCT4 | Fw: ACAGGTCCGCTCTGCAGT<br>Rv: CGTGATGACCCCAGTGGTGA |
| PC | Fw: CATTGCTGCGGGTGTTCC<br>Rv: CCATAGGCCGCCTTGAAGAT |
| TUBA1B | Fw: ACCCACAGTCATTGATGAAGTTCG<br>Rv: CTTGCCAATGGTGTAGTGCC |

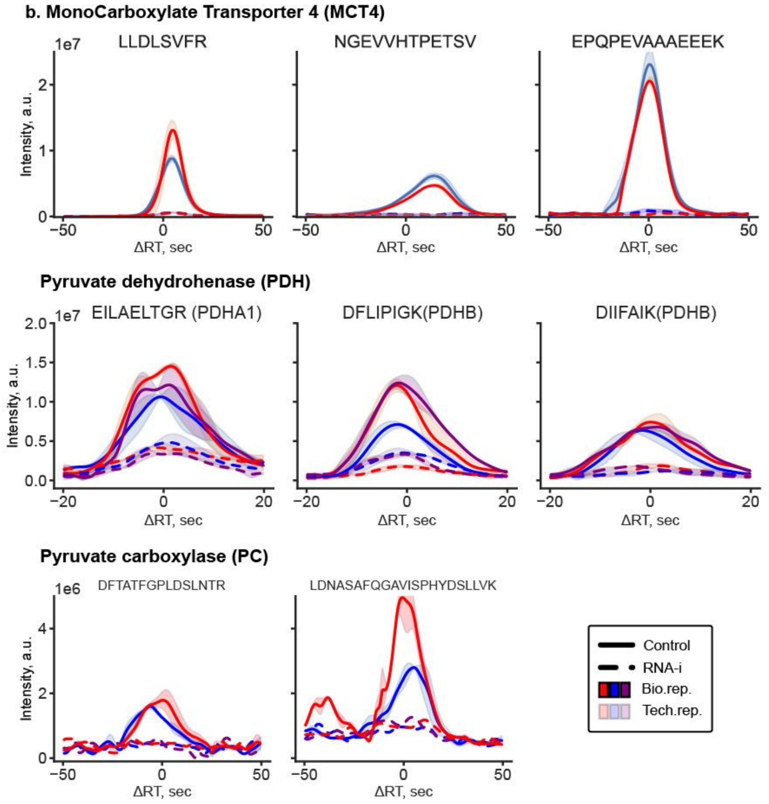


**Extended Data Figure 2. RNA interference a,** shRNA oligonucleotides sequences and relative mRNA expression. **b,** Quality control of RNAi inhibition by targeted proteomics. ±5mDa XICs of selected peptides of targeted proteins in total protein extract of scramble RNA expressed control and shRNA expressed HeLa cells. All chromatograms are aligned and smoothed. Technical replicates are averaged.

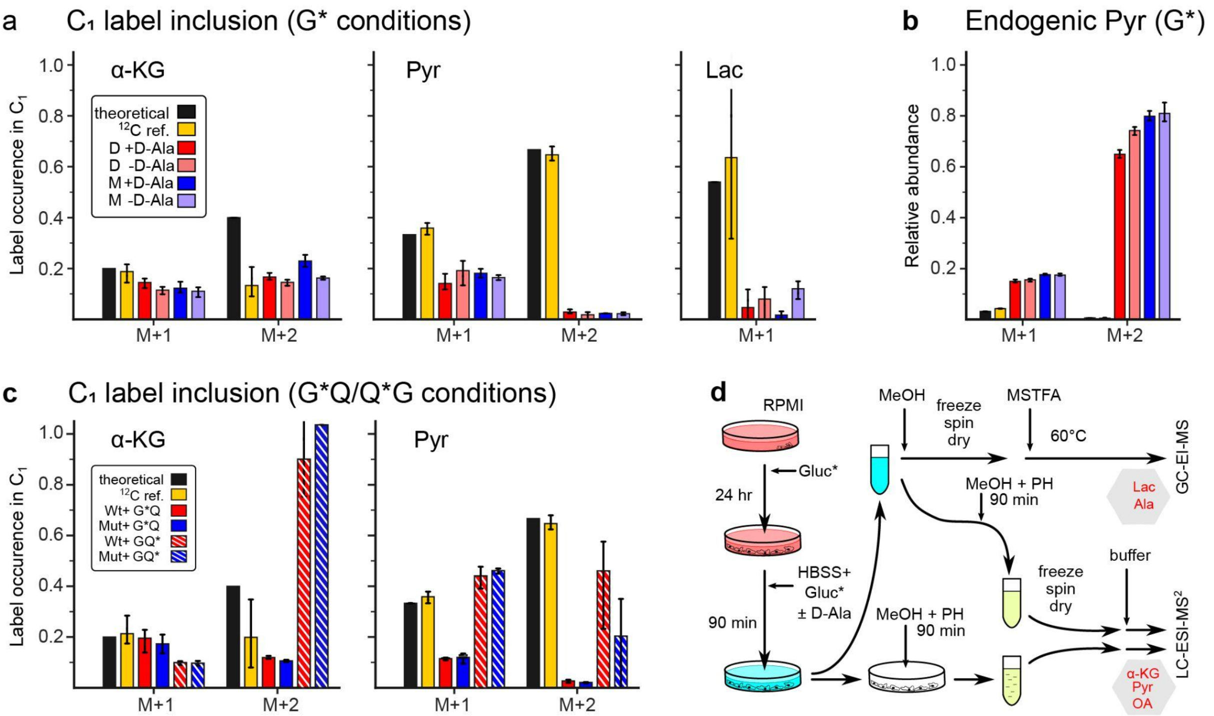


**Extended Data Figure 3. Additional data on isotopologue analysis of metabolic labeling experiments.**

**a,** Occurrence of $^{13}C$ in the first carboxyl atom for an isotopologue of intracellular α-KG, Pyr (measured with LC/MS$^2$) of extracellular Lac (with GC/MS) under conditions with labelled/unlabelled glucose. Missed isotopologues have source chromatogram peaks below signal-to-noise ratio threshold. **b,** Isotopologue abundances for intracellular pyruvate related to M+0 intensity. **c,** Occurrence of $^{13}C$ in the first carboxyl atom for an isotopologue of intracellular α-KG and Pyr (measured with LC/MS$^2$) with parallel labeling of glucose and glutamate. **d,** The design of the experiment. Note that labeled glucose is added twice: to rich RPMI medium (pink) and to pure HBSS medium 90 min before extraction. PH, MSTFA - derivatizing agents. See details in the “Supplementary methods” section.

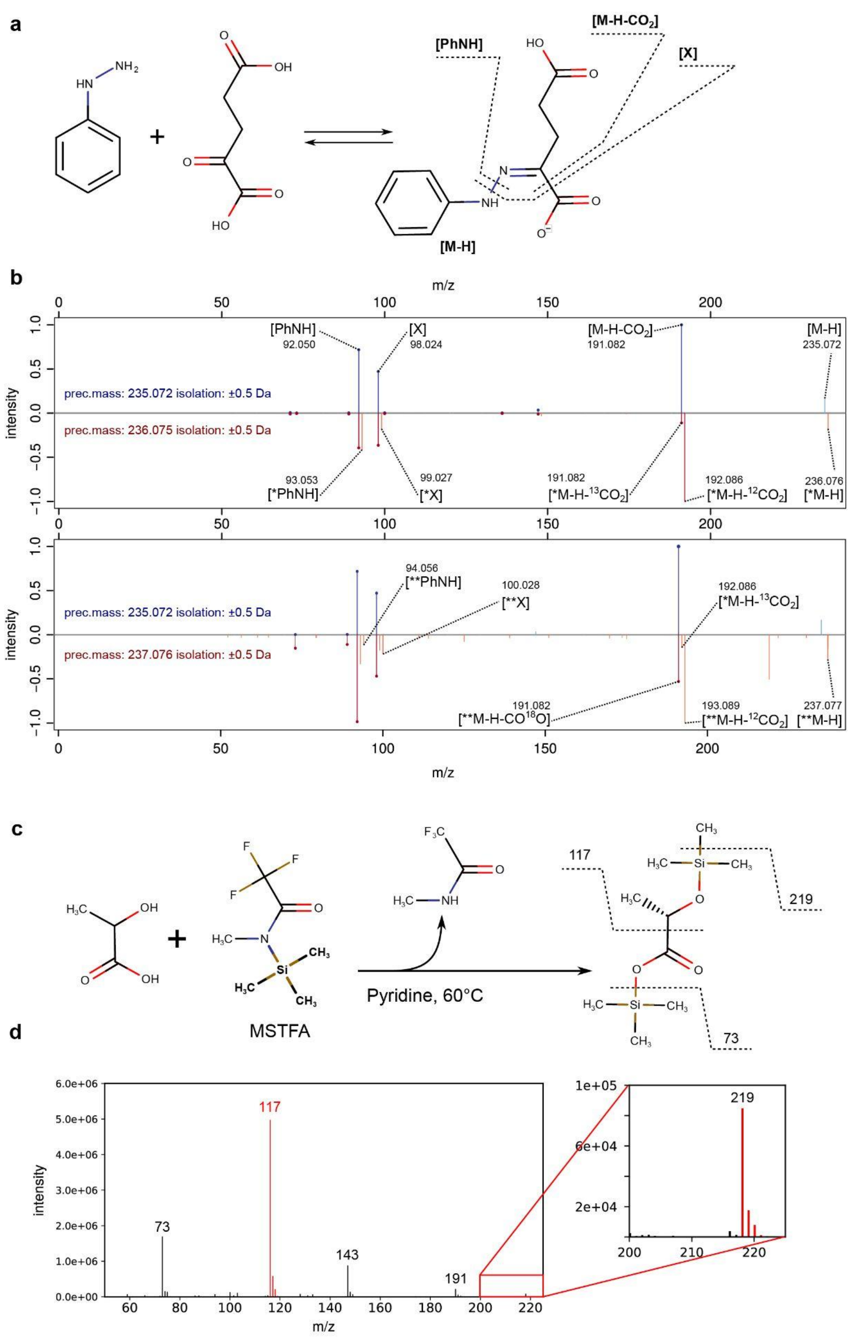


**Extended data Figure 4. Fragmentation analysis of derivatives by LC-MS/MS and GC-MS.** **a,** Derivatization of ketoacids with phenylhydrazine using the example of α-KG. Three identified fragments are shown. **b,** Fragmentation spectra of the following precursor ions: M+1 isotopologue against M+0 (upper) and M+2 against M+0 (lower). MS2 spectrum of monoisotopic precursor ion is above at both panes. Peaks are annotated according to **A**. '*' and '**' superscripts in fragment labels denote fragment's M+1 and M+2 isotopologues respectively. **c,** Derivatization of lactic acid with MSTFA. Three identified fragments are shown. **d,** Scan of the lactic acid peak. The fragments used for analysis are highlighted with red.

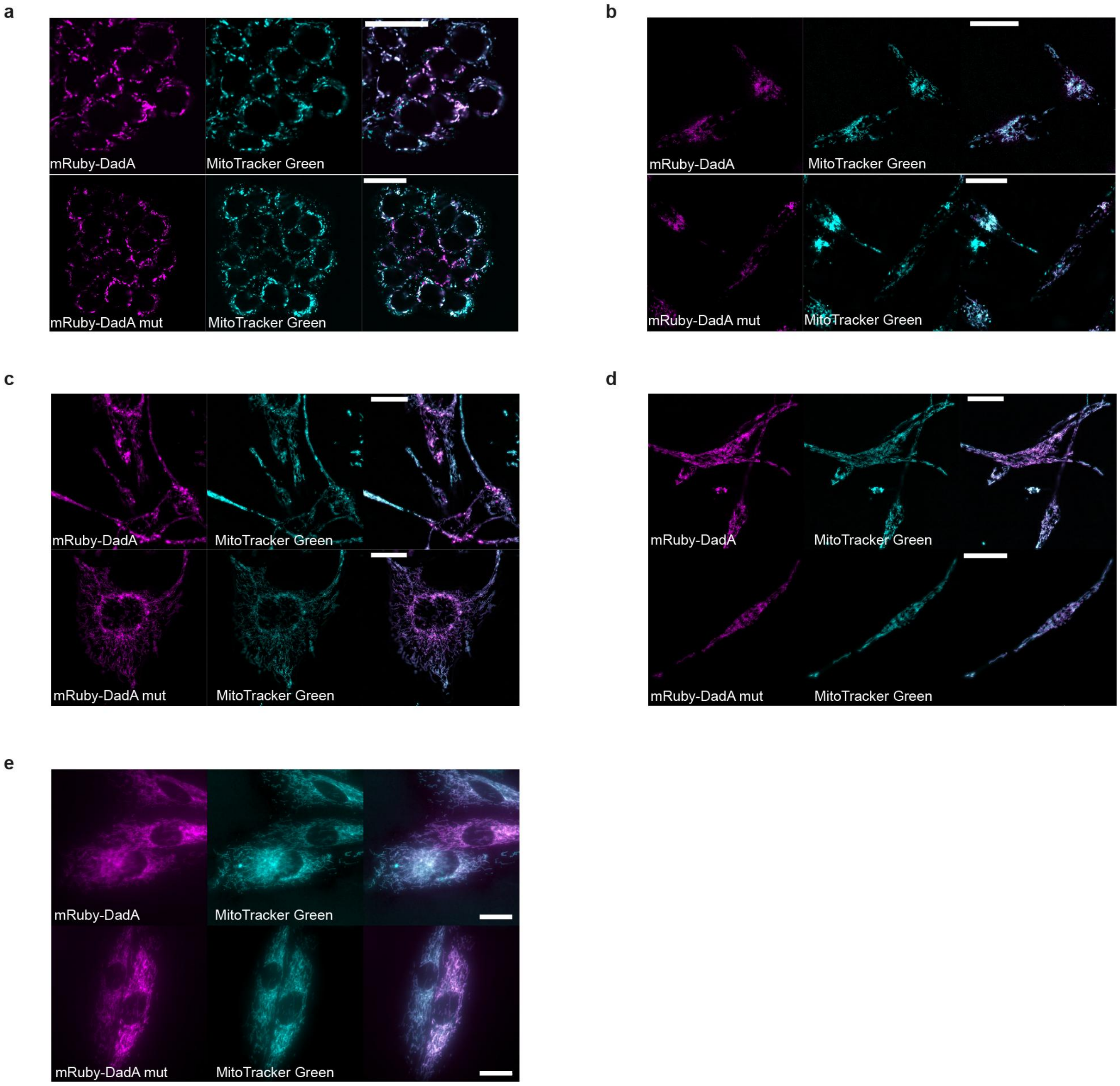


**Extended Data Figure 5. DadA localization in different cell lines.**

Fluorescence image of HT29 (**a**), MDA-MB 231 (**b**), Lu451 (**c**), WM164 (**d**), H9C2 (**e**) cells expressing mito-mRuby-DadA or mito-mRuby-DadA-mut stained with mitoTracker Green. In all fluorescence images, the scale bars represent 25 μm.

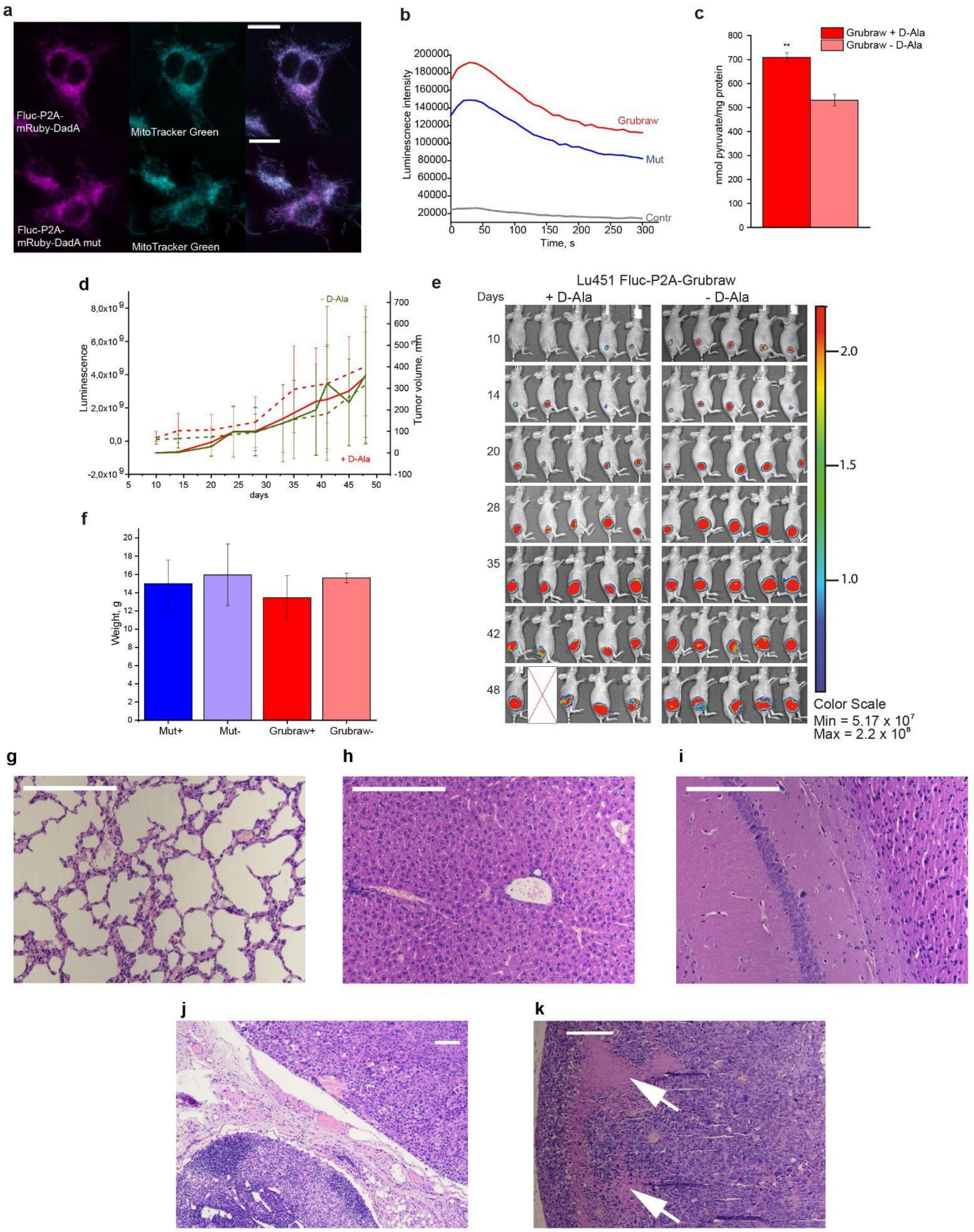


**Extended Data Figure 6. Lu451 Fluc-Grubraw in Nude mice.**

**a,** Fluorescence image of Lu451 cells expressing firefly luciferase (Fluc)-P2A-mito-mRuby-DadA or Fluc-P2A-mito-mRuby-DadA-mut stained with mitoTracker Green. The scale bar represents 25 µm. **b,** Luciferase activity in Lu451 cultures expressing Fluc-P2A-Grubraw (shown in red) and Fluc-P2A-Grubraw-mut (shown in blue). The control sample (shown in gray) contained the same reaction mix without cell culture. The experiment was performed in 1 technical replicate. **c,** The effect of Grubraw activity on extracellular pyruvate in Lu451 cultures expressing Fluc-P2A-Grubraw. **d,** Luminescence (dashed lines) and tumor volume (solid lines) for melanoma xenografts expressing Fluc-P2A-Grubraw-mut. **e,** Bioluminescence in tumors expressing luciferase and mRuby-Gubraw throughout the experiment. **f,** The weight of experimental mice on day 48 after injection. In **c, d, f,** data represent mean values ±SD. The data were compared by t-test. $^{**}p<0.005$. **g – i,** Representative micrographs of experimental mouse lung (**g**), liver (**h**), and brain sections (**i**). **j,** Inguinal lymph node (lower left corner) in the proximity of the subcutaneous tumor (upper right corner). **k,** Representative micrograph of an experimental mouse tumor section. Necrotic regions are shown with white arrows. In **g - k** the scale bars represent 200 μm.

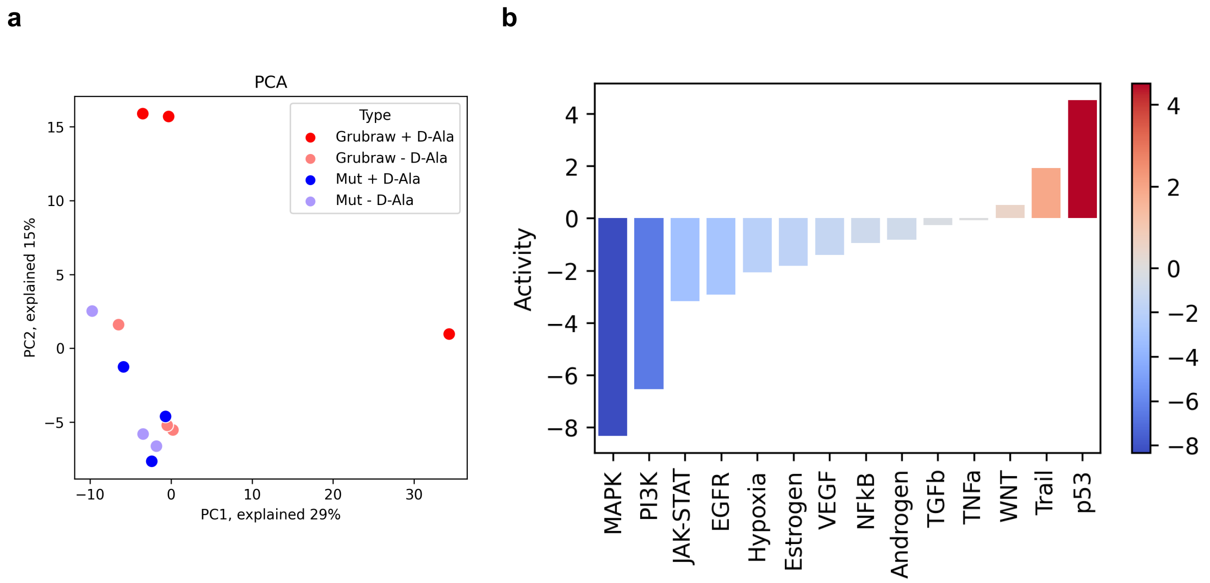


**Extended Data Figure 7. RNASeq analysis on Lu451 cell line.**

**a,** Data heterogeneity based on transcriptomic profiling. Principal component analysis showed similarity between samples with Grubraw construction and no D-Alanin (light red), samples with mutated Grubraw construction with (dark blue) and without (light blue) D-Alanin while samples with Grubraw construction that is activated by D-Ala (dark red) are different from other.

**b,** Pathway activity shift under Grubraw activation. Calculated from transcriptomic data by Progeny. On X axis pathways are shown which are decreased (blue) and increased (red) under Grubraw activation by D-Ala in comparison to all other control samples.

### Supplementary Information

#### Analysis of mass-spectrometry data

In all cases raw LC-MS(2) mass-chromatograms were converted by ThermoRawFileParser[1] into mzML format using its docker version (caetera/thermorawfileparser).

**Targeted proteomics.**

MzML files were analyzed by MSFragger → Philosopher (peptideprohet/filter/proteinprohpet) → IonQuant.[2,3,4] TrEMBL human proteome was used as a search database. We used a ±12ppm window for precursor ions and ±0.05 Da - for fragment ions. Standard variable modifications were used: +15.995 for oxidized methionine, +42.011 for acylated N-end, and +57.021 for carbamidomethylated cysteine. The match-between-run option was turned on in IonQuant runs.

In a treated sample, the concentration of the inhibited protein was too low to be found in PSMs and therefore could not be quantified by the LFQ algorithm. To visualize inhibition efficiency (Extended Data Fig. 2b) we performed alignment of MS1 chromatograms of treated samples to untreated ones using the XCMS library. Peaks were identified with the CentWave algorithm, then retention time was aligned using the OriWarp algorithm.[5] After extracting confident peptides from shotgun runs of control samples, we selected the most abundant ones, identified MS1 peaks and extracted XICs within the 5 mDa window. Chromatograms were then imputed with gaussian background noise at missing points and smoothed by the kernel regression method implemented in the *statsmodels* library.[6]

**Extracting peak intensities from ketoacid data.**

Every sample was recorded twice: one run for LC-MS and one for LC-MS$^2$; each file contains mass-chromatograms for all three SIM ranges. We processed these files with *xcms* library[7] as follows. First, a three-ranged mzML file was split into three mass-chromatograms using *MSnbase* functionality.[8] Then we performed feature detection by CentWave algorithm using preliminarily selected parameters. For MS1-chromatograms we performed peak grouping with the "nearest" method and then aligned chromatograms with linear smoothing (for MS2-chromatograms such alignment is not implemented in *xcms*). Isotopologues of interest were measured as background-corrected intensities of peaks inside a 2 mDa window around precise theoretic m/z values. The latter were calculated using the *enviPat* library.[9] At every stage of processing, aligned XICs were visually controlled. In order to compare samples with different distributions accurately, total compound intensities in MS1 chromatograms were computed as the sum of peaks for *all* isotopologues. Sample R-notebooks with our pipeline have been deposited to a GitHub repository: comcon1/fluxNotebooks.

**Calculating MID of ketoacid using both MS1 and MS2 spectra.**

To extract isotopologue probability distribution for ketoacids, we used the following formula based on the equation of conditional probability:

$$\begin{aligned}[M_{M+1}] &= [Ph_{M+0}|MPh_{M+1}] \cdot [MPh_{M+1}] + [Ph_{M+1}|MPh_{M+2}] \cdot [MPh_{M+2}] \\ [M_{M+2}] &= [Ph_{M+0}|MPh_{M+2}] \cdot [MPh_{M+2}],\end{aligned} \qquad \text{(S1)}$$

where $[M_{M+x}]$ is the occurrence of the M+x isotopologue of the Molecule (without PH), $[MPh_{M+x}]$ is the occurrence of the M+x isotopologue of the derivatized molecule (Molecule + PH)

extracted from LC-MS data, and $[Ph_{M+x}|MPh_{M+y}]$ is the probability of the M+x isotopologue of the PhNH fragment (Extended Data Fig. 3a) in the fragmentation spectrum of the M+y precursor isotopologue.
If the value of M+3 isotopologue occurrence is not negligible, then $[M_{M+1}]$ is still defined by (S1) whereas for other isotopologues the following equation is used:

$$[M_{M+2}] = [Ph_{M+0}|MPh_{M+2}] \cdot [MPh_{M+2}] + [Ph_{M+1}|MPh_{M+3}] \cdot [MPh_{M+3}]$$
$$[M_{M+3}] = [Ph_{M+0}|MPh_{M+3}] \cdot [MPh_{M+3}]. \qquad (S2)$$

Let us explain the origin of the (S1-S2) equations. Ketoacids have one tracer atom in the following cases:
1. the derivative has one tracer atom and it is in the ketoacid
2. the derivative has two tracer atoms and one of them is in PH
3. the derivative has three tracer atoms and two of them are in PH
4. …

Since case 3 has very low probability, we can consider only cases 1 and 2. For the first case, the probability of the derivative having one tracer atom is $[MPh_{M+1}]$. The tracer is in the ketoacid for molecules whose fragment has an M+0 isotopologue of the PhNH fragment, i.e. $[Ph_{M+0}|MPh_{M+1}]$. If we satisfy both conditions, we should multiply them: $[Ph_{M+0}|MPh_{M+1}] \cdot [MPh_{M+1}]$.
The second case is, in turn $[Ph_{M+1}|MPh_{M+2}] \cdot [MPh_{M+2}]$. Adding 1 and 2 we obtain (S1).

**Calculation of the probability of inclusion of the tracer into the C1 position.**

For ketoacid derivatives, the probability of the inclusion of tracer atom into the first position is calculated for every isotopologue from the following equation:

$$[^{13}C_1|M_{M+1}] = [M\text{-}CO2_{M+0}|MPh_{M+1}] / [Ph_{M+0}|MPh_{M+1}]$$
$$[^{13}C_1|M_{M+2}] = [M\text{-}CO2_{M+1}|MPh_{M+2}] / [Ph_{M+0}|MPh_{M+2}] \qquad (S3)$$

.

where $[^{13}C_1|M_{M+x}]$ is the desired probability for M+x isotopologue; $[M\text{-}CO2_{M+x}|MPh_{M+y}]$ is the probability of M+x isotopologue of decarboxylated fragment (Extended Data Fig. 3a) in the fragmentation spectrum of M+y precursor isotopologue.
In equation (S3), the denominator corrects isotopic distribution of the decarboxylated fragment to the natural occurrence of $^{13}C$ isotope in the derivatizing agent.

**Processing GC-MS mass-chromatograms.**

Identification of individual components was carried out using derivatized standards, and by library search (NIST-17). Visual control of GC-MS files in CDF format were also performed using OpenChrom.[10] Metabolite identification was performed using the MassBank and RTX5 FiehnLib[11] libraries downloaded from the Massbank of North America (MoNA) repository.
Files from all experimental samples were loaded by *xcmsSet* function, then peaks were grouped, and retention correction was applied. Next, peaks were grouped again and integrated. Using this method, we extracted data for two fragments of lactate derivative: 219 (full, “F”) and 117 (decarboxylated, “D”) (Extended Data Fig. 3c). These data were firstly corrected for the natural occurrence of tracer in derivatization agent with *IsoCor* package [12]. To derive equations like (S3) for this case, we constructed the following equation system for occurrences of D isotopologues:

$$[D_{M+2}] = [F_{M+2}]\cdot(1 - [^{13}C_1|F_{M+2}]) + [F_{M+3}]$$
$$[D_{M+1}] = [F_{M+1}]\cdot(1 - [^{13}C_1|F_{M+1}]) + [F_{M+2}]\cdot[^{13}C_1|F_{M+2}] \quad (S4)$$

where $[D_{M+x}]$, $[F_{M+x}]$ are the probabilities of isotopologues of decarboxylated and full lactate, and $[^{13}C_1|F_{M+x}]$ is the probability of inclusion of the tracer into the C1 position of lactic acid for M+x lactate isotopologue. Solving this linear system, we obtain the desired probabilities.

**Statistical inference for MID changes.**

The statistical significances of isotopomer occurrence were estimated using the following procedure. First, we gathered all data on every isotopomer occurrence (relating to M+0) of every metabolite pair in all experimental conditions. Next, we computed the t-test pairwise for every row in interesting condition contrasts. Finally, we applied Benjamine-Hochberg correction to the p-values obtained for all isotopomers of all metabolites in one contrast.
